# Microglia reactivity entails microtubule remodeling from acentrosomal to centrosomal arrays

**DOI:** 10.1101/2022.02.21.481355

**Authors:** Maria Rosito, Caterina Sanchini, Giorgio Gosti, Manuela Moreno, Simone De Panfilis, Maria Giubettini, Doriana Debellis, Federico Catalano, Giovanna Peruzzi, Roberto Marotta, Alessia Indrieri, Elvira De Leonibus, Maria Egle De Stefano, Davide Ragozzino, Giancarlo Ruocco, Silvia Di Angelantonio, Francesca Bartolini

## Abstract

Microglia reactivity entails a large-scale remodeling of cellular geometry, but the role of the microtubule cytoskeleton during these changes remains unexplored. Here we show that reactive proinflammatory microglia provide a heretofore unique example of microtubule reorganization from a non-centrosomal array of parallel and stable microtubules to a radial array of more dynamic microtubules. While in the homeostatic state microglia nucleate microtubules at Golgi outposts, proinflammatory signaling induces recruitment of nucleating material nearby the centrosome and inhibition of centrosomal maturation enhances NLRP3 inflammasome activation and secretion of IL-1*β*. Our results demonstrate that a hallmark of microglia reactivity is a striking remodeling of the microtubule cytoskeleton and suggest that pericentrosomal microtubule nucleation may serve as a distinct marker of microglia activation as well as a novel target to modulate cytokine-mediated inflammatory responses in chronic disease and tissue injury.

## INTRODUCTION

Microglia are the brain’s primary innate immune cells. In their homeostatic state in the healthy brain, they exhibit a ramified morphology and continuously patrol the local environment via extension and retraction of highly motile processes ^1^ that act to clear cellular debris, reshape synapses, and provide neurotrophic factors ^2–8^. However, when activated by neuronal inflammation and injury, and in neurodegenerative disorders, microglia exhibit dramatically altered gene expression and morphology, displaying an amoeboid shape ^9–11^. In this pro-inflammatory state, reactive microglia exhibit phagocytic activities that can promote tissue remodeling and if over-activated, are widely thought to contribute to brain damage and neurodegeneration ^2, 12^.

In eukaryotes, changes in cellular symmetry are associated with massive reorganization of both the actin and microtubule (MT) cytoskeletons. While actin and actin-based motor proteins are required for breaking the symmetry in most cells, specification of neuronal polarity depends on MTs and MT associated proteins ^13^. Early studies suggested changes in MT spatial organization and stability also with microglia activation ^14^. Despite these observations, however, only remodeling of the actin cytoskeleton has been extensively studied in microglia, and the role of the MT cytoskeleton in breaking cellular polarity during the transition from an homeostatic to a reactive state has not been explored.

MTs are intrinsically polarized polymers composed of α/β tubulin heterodimers arranged in a head to tail fashion ^15^. They are characterized by a fast growing plus end and a slow growing minus end that in non-neuronal cells is often attached to the centrosome, which acts as the MT organizing center (MTOC). Directional transport is enabled by the structural polarity of MTs, which is recognized by motor proteins that drive transport to either the minus end (dynein) or plus end (most kinesins) (for reviews see ^16, 17^). MTs are generally highly dynamic structures constantly undergoing stochastic transitions from polymerization to depolymerization (catastrophe events) and vice-versa (rescue events), with the two dynamic states exhibiting characteristic rates of growth or shrinkage ^18^. When stabilized, MTs resist disassembly and become substrates of tubulin modifying enzymes that add molecular moieties on either the α- or β-tubulin subunit. The combinatorial nature of these modifications provides a “tubulin code” which controls a variety of functions, including organelle transport and the mechanical properties of the MT lattice (for reviews see ^19–21)^.

MT orientation, density, and post-translational modifications all respond and contribute to breaking cellular symmetry ^22, 23^. Establishment of cell polarity can be achieved through centrosome repositioning or by the formation of non-radial MT arrays, in which MTs are not preferentially nucleated at the centrosome ^24–26^. During this transition, the centrosome typically loses its maximal MT nucleating activity while intracellular membranes or self-organizing assemblies of MT nucleating material distant from the centrosome serve as non-centrosomal MTOCs with members of the calmodulin-regulated spectrin-associated protein (CAMSAP) family often capping and stabilizing released free MT minus ends ^27, 28^. The local stabilization of non-centrosomal MTs and relocation of the MTOCs away from the centrosome and the cell center establish asymmetric MT arrays that are critical to cell differentiation.

We hypothesized that rearrangement of the MT cytoskeleton might be required for the morphological changes that guide microglia transition from surveilling/homeostatic to reactive states. Here we show that proinflammatory microglia engage a unique example of MT transition from a non-centrosomal array of parallel and stable MTs in the homeostatic state to a radial array of more dynamic MTs in which all MT minus ends are anchored to a pericentrosomal region. We further find that in the homeostatic state, Golgi outposts are sites of non-centrosomal MT nucleation, and that a pro-inflammatory challenge leads to the recruitment of pericentriolar material (PCM) to the centrosome. To investigate the regulatory role of this transition we inhibited the master modulator of MTOC assembly Polo-like kinase 4 (PLK4) and found that failure to mature *de novo* formed pericentrosomal MTOCs increases the number of small diameter extracellular vesicles (EVs) and selectively enhances IL-1*β* secretion. Our results unveil the unique rearrangement of the MT cytoskeleton in proinflammatory microglia and indicate that pericentriolar material re-localization and assembly can alone limit the release of IL-1*β*.

## MATERIALS AND METHODS

### Primary murine microglia culture and treatment

Primary cortical glial cells were prepared from 0- to 2-d-old mice as previously described ^29^. Briefly, cerebral cortices were chopped and digested in 30 U/ml papain for 40 min at 37 °C followed by gentle trituration. The dissociated cells were washed, suspended in Dulbecco′s Modified Eagle′s Medium (DMEM, Sigma-Aldrich by Merck KGaA, Darmstadt, Germany) with 10% FBS (Gibco by Life Technologies, Carlsbad, CA, USA) and 2 mM L-glutamine and plated at a density of 9–10 x 105 in 175 cm^2^ cell culture flasks. At confluence (10 –12 DIV), glial cells were shaken for 2 h at 37 °C to detach and collect microglial cells. These procedures gave an almost pure (<1% astrocyte contamination) microglial cell population. Microglia cells were plated at a density of 7x103/cm^2^ (to prevent cell contact activation) in astrocytes conditioned medium/DMEM 2,5% FBS (1:1). The day after plating microglia cells were treated for 48 h with IFNγ (20 ng/ml) and LPS (100 ng/ml) or with IL-4 (20 ng/ml) to obtain the pro-inflammatory or anti-inflammatory phenotype, respectively. To disassemble MTs, homeostatic microglia were treated with 2 µM nocodazole (Sigma-Aldrich) added to the culture medium for 1 h at 37 °C. Samples were then kept on ice and washed 5x times with ice-cold medium. MTs were allowed to regrow in conditioned medium without nocodazole for 15 min and 120 min at 37 °C. Right before fixation, free tubulin was rapidly extracted using a MT-preserving extraction buffer (60 mM PIPES, 25 mM HEPES, 10 mM EGTA, 2 mM MgCl2, 0.1% saponin, pH 6.9, ^30^) for 20 sec at 37 °C. Cells were subsequently fixed with methanol at -20 °C for 4 min and processed for immunofluorescence staining. To stabilize MTs, microglia cells were treated for 24 h with 1 nM and 5 nM Taxol (Sigma-Aldrich) alone or together with IFNγ (20 ng/ml) and LPS (100 ng/ml). To inhibit PLK4, microglia cells were pre-treated with 125 nM Centrinone (Tocris Bioscience, Bristol, UK) for 12 h prior to stimulation for 48 h with IFNγ (20 ng/ml) and LPS (100 ng/ml).

### Immunofluorescence staining on fixed cells

Methanol fixation at -20 °C was elected for preserving an intact MT cytoskeleton: culture medium was removed and cells were fixed with pre-cooled 100% methanol at -20°C for 4 min prior to re- hydration with Phosphate-buffered saline (PBS, 0.01 M phosphate buffer, 0.0027 M potassium chloride and 0.137 M sodium chloride, pH 7.4, at 25 °C, Sigma-Aldrich) for at least 30 min at RT To preserve membrane associated components, cells were fixed with 4% paraformaldehyde (PFA)/PBS for 15 min at RT and then washed with PBS. When PFA fixed, cells were permeabilized with 0.1% Triton X-100/PBS for 1 to 3 min. After 2 washes in PBS, cells were blocked with 3% bovine serum albumin (BSA, Sigma-Aldrich) in PBS for 1 h at RT Primary antibodies (Rabbit Camsap1L1, Novus Biologicals, Englewood, CO, USA, 1:200; rabbit γ-tubulin, Invitrogen, Waltham, MA, USA, 1: 5000; mouse γ tubulin, Sigma-Aldrich, 1:1000; rat Tyrosinated tubulin YL 1/2, Merck-Millipore, 1:1000; mouse Acetylated tubulin clone 6-11B-1, Sigma-Aldrich, 1:1000; mouse α-tubulin clone DM1A, Sigma-Aldrich, 1:500; rabbit Detyrosinated α- tubulin, Merck-Millipore, 1:1000; mouse EB1, BD Biosciences, San Jose, CA, USA 1:100; rabbit Pericentrin, Abcam, Cambridge, UK, 1:1500; mouse Centrin3, Abnova, Taipei City, Taiwan, 1:100; mouse GM130, BD Biosciences, 1:600; rabbit IBA-1, FujiFilm Wako, Richmond, VA, 1:300; Atto 488 Phalloidin, Sigma-Aldrich, 1:50) were incubated in 1.5% BSA in PBS for 2 h (RT) or overnight (+4°C). Cells were then extensively washed and stained with fluorophore-conjugated secondary antibodies in PBS (Alexa Fluor 488 goat anti-mouse, 488 goat anti-rat, 488 goat anti-rabbit, 594 goat anti-mouse, 647 goat anti-rabbit, Invitrogen; CF 594 goat anti-rat, Sigma-Aldrich; 1:500) and Hoechst (Sigma-Aldrich) for nuclei visualization for 1 h at RT prior to wash and mounting using Ibidi Mounting Medium.

### MT dynamics assay

For live fluorescence imaging to measure MT dynamics, cells were incubated with 100 nM SiR-Tubulin (SpiroChrome, Stein-am-Rhein, Switzerland) at 37 °C for 30 min and washed with conditioned medium prior to visualization to improve signal to noise ratio. 10 µM Verapamil was added to inhibit efflux pumps and improve labeling. Live wide-field fluorescence imaging of SiRTub-labeled MTs was performed on a Olympus IX73 microscope, LDI laser source and CoolSNAP Myo camera, 4.54 µm pixels (Photometrics, Tucson, AZ, USA) with a built-in incubator, maintaining the temperature at 37 °C during recordings. Acquisitions were performed for 4 min (1 frame/4 sec) with a UPLSXAPO100x/1.45 oil objective and then analyzed with ImageJ software (see Image preparation and analysis).

### Animals

All procedures performed using laboratory animals were in accordance with the Italian and European guidelines and were approved by the Italian Ministry of Health in accordance with the guidelines on the ethical use of animals from the European Communities Council Directive of September 20, 2010 (2010/63/UE). All efforts were made to minimize suffering and number of animals used. Mice were housed in standard cages in a group of a maximum of 5 animals, with light–dark cycles of 12 h at 22±2 °C. Wild type C57BL-6 male and pregnant mice were purchased from Charles River and pups (P0-P2) were used to obtain primary glial cultures. Cx3cr1^gfp/gfp^ male mice were purchased from The Jackson Laboratory company (B6.129P2(Cg)-Cx3cr1tm1Litt/J); the colony was established in our animal facility, and progenitors were bred to C57BL6J to obtain cx3cr1^gfp/+^ mice as we previously reported ^31^.

### Intravitreal injection and EIU

Adult C57BL6/J mice were intravitreally injected with sterile PBS (vehicle) or 5 ng/µl LPS from E. Coli (O55:B5, Sigma Aldrich). Intravitreal injection of LPS has been previously reported as a model of endotoxin induced uveitis (EIU) activating microglia in the retina ^32–36^. Animals were anaesthetized with 100 mg/kg methadomidine and 0.25 mg/kg ketamine. Pupils were dilated using 1% tropicamide and 2.5% phenylephrine (Chauvin, Essex, UK) and a small guide hole was made under the limbus with a 30G needle. The eye was gently massaged with a cotton swab to remove a portion of the vitreous to avoid a post-injection reflux of vitreous and/or drug solution. Then, 1 µl of vehicle or LPS solution was intravitreally injected through the initial hole using a 34G Hamilton syringe.

### Immunofluorescence staining on retinal tissue

Cx3cr1^gfp/+^ control mice were sacrificed at P70. CTRL (sham) and LPS intravitreally injected adult C57BL6/J mice were sacrificed 20 h after the injection procedure. Eyes were removed and kept in 4% PFA solution overnight. Eyes were then cryoprotected in 30% sucrose and, after precipitation, frozen in isopentane prior to storage at -80 °C. Frozen eyes were cut in 50-μm-thick sections with a Leica cryostat and processed for immunofluorescence staining as published ^37^. Briefly, slices were immersed for 30 min in a boiling 1 mM EDTA solution (pH = 8.0) for antigen retrieval, then incubated with blocking solution (0.1% Triton X-100, 3% BSA and 0.05% Tween-20 in PBS) for 1 h at RT. Sections were incubated with primary antibodies (Iba1, FujiFilm Wako, 1:500; γ-tubulin, clone GTU-88, Sigma-Aldrich, 1:500; GM130, BD bioscience, 1:500) in diluted blocking solution overnight at 4 °C and 1 h at RT with fluorophore-conjugated secondary antibodies (Alexa Fluor 488 goat anti-rabbit, 594 goat anti-mouse) and Hoechst for nuclei visualization. The sections were mounted with anti-fade mounting medium (Invitrogen).

### Confocal Spinning Disk and Structured Illumination (SIM) microscopy

For fluorescence imaging of fixed samples, images were collected with spinning disk confocal microscopy on a Nikon Eclipse Ti equipped with X-Light V2 spinning disk (CrestOptics, Rome, Italy), combined with a VCS (Video Confocal Super resolution) module (CrestOptics) based on structured illumination, and a LDI laser source (89 North, Williston, VT, USA) and Prime BSI Scientific CMOS (sCMOS) camera, 6.5 µm pixels (Photometrics) or a CoolSNAP Myo camera, 4.54 µm pixels (Photometrics), with a 10x/0.25 NA Plan E air objective, 40x/0.75 PlanApo l air objective, a 60x/1.4 PlanApo l oil objective and a 100x/1.45 Plan E oil objective. The used Z step size was 0.2 µm for spinning disk and 0.1 µm for VCS. In order to achieve super-resolution, raw data obtained by the VCS module have been processed with a modified version of the joint Richardson-Lucy (jRL) algorithm ^38–40^, where the out of focus contribution of the signal has been explicitly added in the image formation model used in the jRL algorithm, and evaluated as a pixel-wise linear “scaled subtraction” ^41^ of the raw signal. Retinal sections images were acquired on an Olympus IX73 microscope equipped with X-Light V3 spinning disk (CrestOptics), LDI laser source and a Prime BSI Scientific CMOS (sCMOS), 6.5 µm pixels (Photometrics) with a UPLSXAPO100x/1.45 oil objective. All the images were acquired by using Metamorph software version 7.10.2. (Molecular Devices, Wokingham, UK) and then analyzed with ImageJ software (see Image preparation and analysis).

### Image preparation and analysis

For image preparation, we used the open-source software ImageJ ^42^ for adjustments of levels and contrast, maximum intensity projections, and thresholding signals for fluorescence intensity analysis.

#### Radial profile analysis

For tyrosinated α-tubulin, CAMSAP2 and γ-tubulin distribution analysis, microglia cells were fixed in methanol at -20 °C for 4 min or PFA 4% for 15 min and then stained with an anti-tyrosinated tubulin, CAMSAP2 or γ-tubulin antibody according to the immunofluorescence protocol, and Hoechst for nuclei visualization. Images obtained by confocal microscopy were analyzed with ImageJ to identify the coordinates of the center of the nucleus in each cell and to generate single-cell masks based on the morphology of each cell. A Python script (see Supplementals) was written to apply an Otsu threshold to the images ^43^ and to perform a radial scanning of fluorescence values, starting from the center of the nucleus of each cell, with a resolution of 0.065 μm. Maximum value radial profile was defined as the maximum fluorescence intensity (a.u.) for each concentric circle with an increasing distance from the nucleus center. For each analyzed microglia cell the radial profile of the maximum value of fluorescence intensity (a.u.) was computed and plotted. Plots were smoothed with a resolution of 0.5 μm. All data points were exported into a Microsoft Excel 2010 compatible format. In CAMSAP2 analysis, only cytoplasmic staining was analyzed. Curve fit was performed using a single exponential decay function on GraphPad Prism 9.0 (Y=(Y0 - Plateau)*exp(-K*X) + Plateau).

MT dynamics analysis. Analysis of MT dynamics was performed by tracing the lengths of the MTs via the “freehand line” tracing tool in ImageJ. Changes in length between successive frames were exported into an Excel sheet to determine the growth, shortening and pause events for each MT. Only changes >0.5 µm were considered growth or shortening events ^44, 45^. MT dynamics parameters were defined as follows: growth/shrinkage rate: distance (µm) covered in growth or shrinkage per second; % pause/growth/shrinkage: number of frames in pause/growth/shrinkage divided total number of frames X 100; catastrophe/rescue frequency (sec-1): number of catastrophe or rescue events divided by the product of the time of analysis and the percentage of growth or shrinkage; MT dynamicity: the sum of total length in growth and shortening divided by the time of analysis.

#### In vitro cell morphology analysis

Cell morphology analysis was performed using a quantitative measurement of cell area; cell solidity is expressed as the ratio between cell area and convex area. Measurements were obtained with the Particle Analysis tool and images were processed with ImageJ.

#### Extracellular vesicle analysis

For the statistical analysis of EV blebbing from the surface of microglia, 20 microglia cells per sample, collected in four different areas of the support, were randomly selected and scanned to count and measure the visualized vesicles. The ImageJ software was used to count and measure the vesicle major axis.

#### Immunofluorescence signal quantification

For immunofluorescence signal quantification, cells were selected based on the representative morphology: ramified for homeostatic, ameboid for pro-inflammatory and bipolar for anti-inflammatory states. Detyr/Tyr tubulin ratio and Acetyl/Tyr tubulin ratio were calculated from the mean gray values of the respective immunofluorescence signals, obtained from sum slices z-projections of 15 confocal planes after background subtraction (calculated as mean gray value of three circle background areas). EB1 anterograde or retrograde comets were defined from the EB1 fluorescence signal gradient from single plane images, measured with the “plot profile” tool of ImageJ. For Golgi stacks analysis, GM130 maximum intensity z-projection immunofluorescence images were uniformly thresholded on ImageJ by setting the same minimum values (‘Default’ threshold) to identify single Golgi stacks; a single Golgi stack was defined as a non-round object (roundness < 0.9) with a major axis length > 0.5 µm. For GM130- γ-tubulin co-staining analysis, GM130 and tubulin signals from max intensity z-projections were uniformly processed among different images increasing the ‘brightness’ and ‘contrast’ parameters by the same percentage. *γ*-tubulin signal over cell area was calculated as percentage of cell area covered by γ tubulin signal; γ-tubulin signal threshold was uniformly applied on MetaMorph analysis software by setting the same minimum values to all images. Two or more distinct γ-tubulin^+^ puncta were identified by counting the peaks of fluorescence intensity on a linescan drawn through the centroid of each puncta using the free-hand tool on ImageJ; The Find Peaks ImageJ plugin was used to identify the peaks by setting the minimum peak amplitude value at 100 grey values. Puncta were identified in a 132x132 pixels pericentriolar region after uniformly thresholding max intensity z-projections (‘Default’ threshold); integrated density was calculated in ImageJ as mean gray value*thresholded area. Pericentrin and Centrin-3- γ-tubulin co-localization analysis was performed by defining Pericentrin and Centrin-3^+^ puncta as described above for γ-tubulin^+^ puncta.

#### Retinal microglia cell skeleton analysis

Morphology of microglia cells in retinal sections was analyzed on max intensity z-projections; only entirely visible cells inside the acquisition field were analyzed; cells were isolated and then skeletonized on binary images, using the dedicated ImageJ plug-in; branches, endpoints and junction number was calculated from the skeletonized image.

### Real time PCR

RNA was extracted from microglia cells with the Quick RNA MiniPrep (Zymo Research, Freiburg, DE) and retrotranscribed with iScript Reverse Transcription Supermix for Real-time PCR (RT-PCR) (Bio-Rad, Hercules, CA, USA). RT-PCR was carried out using Sybr Green (Bio-Rad) according to the manufacturer’s instructions. The PCR protocol consisted of 40 cycles of denaturation at 95 ◦C for 30 s and annealing/extension at 60 ◦C for 30 s. For quantification, the comparative Threshold Cycle (Ct) method was used. The Ct values from each gene were normalized to the Ct value of GAPDH in the same RNA samples. Relative quantification was performed using the 2−ΔΔCt method ^46^ and expressed as fold change in arbitrary values. The primers were used as below: GAPDH forward TCGTCCCGTAGACAAAATGG; GAPDH reverse TTGAGGTCAATGAAGGGGTC; Ym1 forward CAGGTCTGGCAATTCTTCTGAA; Ym1 reverse GTCTTGCTCATGTGTGTAAGTGA; Fizz1 forward CCAATCCAGCTAACTATCCCTCC; Fizz1 reverse ACCCAGTAGCAGTCATCCCA; Tnfα forward GTGGAACTGGCAGAAGAG; Tnfα reverse CCATAGAACTGATGAGAGG; IL1*β* forward GCAACTGTTCCTGAACTCAACT; IL1 *β* reverse ATCTTTTGGGGTCCGTCAACT; iNOS forward ACATCGACCCGTCCACAGTAT; iNOS reverse CAGAGGGGTAGGCTTGTCTC.

### SEM analysis

Samples were fixed in a solution of 1.5% Glutaraldehyde in 0.1 M Cacodylate buffer for 2 h at RT and post-fixed in 1% osmium tetroxide in Milli q (MQ) H2O for 2 h. After several washes in MQ H2O, the samples were subsequently dehydrated in rising concentrations of ethanol in H2O solutions (from 30% to 100%), 1:1 ethanol:hexamethyldisilazane (HMDS, Sigma-Aldrich) and 100% HMDS and dried overnight in air. Finally, the samples were sputtered with a 10 nm gold layer and analyzed using a JEOL JSM-6490LA Scanning Electron Microscope (SEM) operating at 10 KV of accelerating voltage.

### Western blot

Cells were lysed in Laemmli sample buffer and boiled at 95 °C for 5 min. Proteins were separated by 4-12% Bis-Tris gel (Invitrogen) and transferred onto nitrocellulose membrane. After blocking in 5% milk/TBS (Tris 20 mM, NaCl 150 mM), membranes were incubated with primary antibodies at 4 °C overnight prior to 1 h incubation with secondary antibodies and signal detected using a commercial chemiluminescent assay (Immun-Star WesternC Kit; Bio-Rad). Image acquisition was performed with ChemiDoc MP imaging system (Bio-Rad) and densitometric analysis was performed with Quantity One software (Bio-Rad).

### Cell cycle assay

Microglia cells were collected by following trypsin treatment. Cells were rinsed twice with phosphate buffered saline (PBS pH 7.4) and collected by centrifugation. Pellets were resuspended in ice cold 70% ethanol and stored at 4 °C for 1 h. Cells were collected by centrifugation, rinsed twice in PBS and resuspended in 20 μg/ml propidium iodide (PI) in PBS with 50 μg/ml RNase A for a minimum of 30 min. After PI incubation, Flow cytometry analysis of DNA content was performed as reported and analyzed using a BD LSRFortessa (BD Biosciences). The percentage of cells in different phases of the cell cycle was determined using the FlowJo V10.7.1 computer software (TreeStar, Ashland, OR, USA). At least 10.000 events for each sample were acquired.

### Statistical analysis

The n number for each experiment and details of statistical analyses are described in the figure legends or main text. Data are reported as mean ± SEM; when not normally distributed, data are reported as median ± interquartile range. Origin 6 and GraphPad Prism 9 software were used for statistical analysis. Normality tests were performed with Prism 9 and nonparametric tests were used when appropriate. Significant differences are indicated in the figures by * p <0.05, ** p <0.01, *** p <0.001. Notable non-significant differences are indicated in the figures by ns.

## RESULTS

### Homeostatic, pro-inflammatory and anti-inflammatory primary microglia differ in MT distribution, stability and dynamic behaviour

To investigate the organization of the MT cytoskeleton in homeostatic and reactive microglia, we used primary mouse microglia cultures in which the presence of ramified cells was maintained by growth factors secreted by astrocytes ^47^. With this approach we prepared a nearly pure population of primary microglia comprised by 99% of Iba1 positive cells. To steer microglia towards different reactivity states such as a pro-inflammatory or an alternatively polarized microglia state (defined as anti-inflammatory), cells were challenged with either LPS-IFNγ (100 ng/ml – 20 ng/ml 48 h for pro-inflammatory) or IL-4 (20 ng/ml, 48 h for anti-inflammatory), and measured for the expression of their signature activation genes (Figure S1A). As revealed by Iba1 staining (Figure S1B), polarized microglia underwent dramatic morphological changes. We classified cell morphology as ramified (*≥*3 ramifications), ameboid or bipolar based on number of cellular processes, cell area, and solidity, a measure of cell shape complexity (Figure S1B-D). Analysis of morphology distribution under homeostatic, pro-inflammatory and anti-inflammatory conditions revealed that ramified cells were enriched in untreated microglia (35 ± 3%) (Figure S1B and S1D), while ameboid cells represented a large majority after pro-inflammatory stimulation (82 ± 3%) (Figure S1B and S1D). Conversely, when cells were challenged with an anti-inflammatory stimulus, microglia mostly acquired a unipolar or bipolar rod-shape morphology (54 ± 3%) characterized by the presence of a lamellipodium and a trailing edge, or uropod (Figure S1B and S1D). To further detail the structural changes associated with reactive microglia states *via* single-cell analyses, we chose to select for comparison only the most representative morphology of each *in vitro* phenotype (ramified for homeostatic, amoeboid for pro-inflammatory and bipolar for anti-inflammatory microglia).

We employed scanning electron microscopy (SEM) and confocal microscopy to identify defined ultrastructural elements typical of each functional state (Figure S1E). Homeostatic microglia exhibited many branched processes extending outward from the cell body and multiple filopodia-like structures (Figure S1E) that were also positive for phalloidin staining (Figure S1H). Upon pro-inflammatory challenge, microglia retracted most of their processes and acquired a flattened and round morphology (Figure S1E and S1H). SEM imaging further revealed that pro-inflammatory microglia displayed numerous tethered extracellular vesicles (EVs) blebbing from the cell surface (Figure S1F). Analysis of EV diameter showed a bell-shaped distribution of size, ranging from 250 to 650 nm (Figure S1G), consistent with microglia-shedded microvesicles ^48^. Anti-inflammatory microglia were characterized by extensive membrane ruffling at both uropod and leading edge, which appeared as sheet-like structures on the dorsal cell surface (Figure S1E and S1H).

We began to analyze the MT cytoskeleton in each microglia functional state by immunofluorescence staining of tyrosinated *α*-tubulin (Tyr tub), a bulk tubulin marker labeling the entire MT network. MTs appeared to be packed in a parallel fashion in all the cellular branches extending from the cell body in both homeostatic and anti-inflammatory microglia (Figure 1A). However, MTs distributed radially from a perinuclear region in pro-inflammatory microglia (Fig. 1A). Radial profiling of Tyr tub fluorescence intensity (Figure 1A), a measure of the distribution of tubulin signal that is independent of cell shape, confirmed that MT staining was uniformly distributed along the entire cell profile in homeostatic and anti-inflammatory microglia (Figure 1B), while in pro-inflammatory cells, Tyr tub signal rapidly decayed at increasing distances from a perinuclear region (Figure 1B; exponential decay constant k_pro-inf_= 0.046 ± 0.001; k_homeo_= 0.013 ± 0.002; k_anti-inf_= 0.015 ± 0.002).

**Figure 1.**
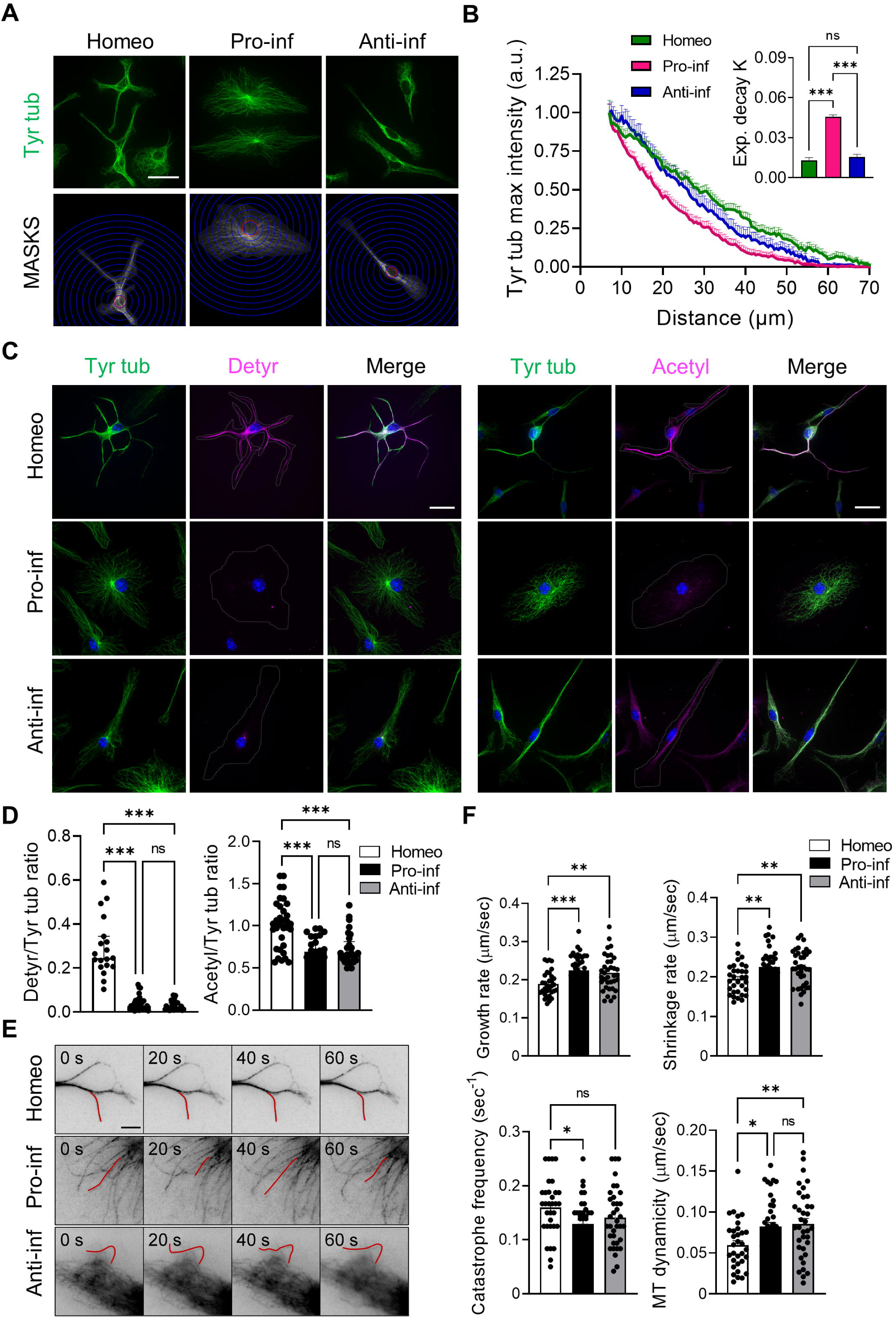
Homeostatic, pro-inflammatory and anti-inflammatory primary microglia differ in MT distribution, stability and dynamic behavior. (**A**) Representative images of tyrosinated α-tubulin (Tyr tub, green) staining in homeostatic (Homeo), pro-inflammatory (Pro-inf) and anti-inflammatory (Anti-inf) microglia (*top*; scale bar: 20 µm) and corresponding masks (MASKS) used for the analysis (*bottom*), with radial scale (unit = 5 μm; blue circles) centered at the centroid of the cell nucleus. Red circle indicates the radius corresponding to the largest intensity value. (**B**) Plot showing maximum fluorescence intensity values of Tyr tub vs the radial distance from cell nucleus, obtained with radial profiling, in Homeo (n = 38, green), Pro-inf (n = 31, magenta) and Anti-inf (n = 38, blue) microglia. Curve fit was performed using a single exponential decay function. *Insert:* bar chart reporting the exponential decay constant values (K) for each condition (values are expressed as mean ± SEM from 4 independent experiments; *** p <0.001, One-way ANOVA – Tukey’s multiple comparison test). Note faster decay of Tyr tub signal in Pro-inf microglia. (**C**) Representative immunofluorescence images of Homeo, Pro-inf e Anti-inf microglia: *left*, co-staining of Tyr tub (green) and de- tyrosinated tubulin (Detyr, magenta) (scale bar: 20 µm. Hoechst for nuclei visualization, blue); *right*, co-staining of Tyr tub (green) and acetylated tubulin (Acetyl, magenta) (scale bar: 20 µm. Hoechst for nuclei visualization, blue). Cell outlines are indicated by white dashed line. (**D**) Scatter dot plots showing immunofluorescence signal quantification of detyrosinated/ tyrosinated (Detyr/Tyr) tub ratio (*left*, Homeo n = 19, Pro-inf n = 32, Anti-inf n = 28 cells from 3 independent experiments) and acetylated/tyrosinated (Acetyl/Tyr) tubulin ratio (*right*; Homeo n = 33, Pro-inf n = 25, Anti-inf n = 29 cells from 3 independent experiments). Values are expressed as median ± interquartile range; *** p <0.001; Kruskall-Wallis - Dunn’s multiple comparisons test. Note that Pro- and Anti-inf microglia have reduced tubulin PTM levels with respect to Homeo cells. (**E**) Representative inverted contrast widefield frames from time lapse acquisitions of SiRtubulin in Homeo, Pro-inf and Anti-inf microglia at 4 different timepoints (0 s, 20 s, 40 s, 60 s). Red lines highlight MT length changes ≥0.5 μm between frames. Scale bar: 5 μm. (**F**) Scatter dot plots representing growth rate (*top*, *left*), shrinkage rate (*top*, *right*), catastrophe frequency (*bottom*, *left*) and MT dynamicity (*bottom*, *right*) in Homeo, Pro-inf and Anti-inf microglia. Values are expressed as mean ± SEM (Homeo n = 30, Pro-inf n = 27 and Anti-inf n = 29 cells from 4 independent experiments). *** p <0.001; ** p <0.01; * p <0.05. One-way ANOVA - Dunnett’s multiple comparison test.

To evaluate whether microglial MTs differed in stability according to their reactive state, we analyzed levels and distribution of detyrosinated and acetylated tubulins, two independent tubulin post-translational modifications (PTMs) associated with MT longevity ^19, 49^. Semiquantitative immunofluorescence analyses revealed that homeostatic cells had the highest level of both detyrosinated (Figure 1C, left) and acetylated (Figure 1C, right) tubulin compared to pro-inflammatory or anti-inflammatory microglia (De-tyr/Tyr tub: 0.30± 0.03; 0.04 ± 0.01; 0.03 ± 0.01, Acetyl/Tyr tub: 1.02 ± 0.05; 0.69 ± 0.04; 0.72 ± 0.04 in homeostatic, pro-inflammatory and anti-inflammatory microglia respectively; Figure 1D), suggesting that homeostatic microglia display more stable MTs.

We quantified the behavior of SiR-Tubulin labeled MTs ^50^ in shallow peripheral sections of the cell to measure MT plus end dynamics using time lapse wide field fluorescence microcopy (Figure 1E). No change was observed among the three different microglia states in rescue frequency (frequency of transitions from shrinkage to growth) or the fraction of time spent in pausing or shrinkage. However, while pro-inflammatory and anti-inflammatory microglia MTs exhibited a moderate yet significative drop in catastrophe frequency (frequency of transitions from growth to shrinkage), they also significantly enhanced their growth rates and acquired a nearly 2.5 fold increase in rates of shrinkage, resulting in an overall net rise in MT dynamicity compared to MTs of homeostatic cells (0.06 ± 0.01; 0.08 ± 0.01; 0.09 ± 0.01 in homeostatic, pro-inflammatory and anti-inflammatory microglia respectively; Figure 1F, Table 1, Movies S1-3). In summary, and consistent with our analysis of tubulin PTMs, these data suggest that microglia acquisition of pro- and anti-inflammatory phenotypes is characterized by loss of MT stability and a marked increase in MT dynamics.

**Table 1.** MT dynamicity parameters in homeostatic and activated microglia. Table reporting parameters of MT dynamics obtained from wide field fluorescence time lapse analysis of SiRTubulin in homeostatic (Homeo), pro-inflammatory (Pro-inf) and anti-inflammatory (Anti-inf) microglia. Values are expressed as mean ± SEM from Homeo n = 32/30 MTs/cells, Pro- inf n = 40/27 MTs/cells and Anti-inf n = 36/29 MTs/cells arising from 4 independent experiments. *** p <0.001; ** p <0.01; * p <0.05. One-way ANOVA - Dunnett’s multiple comparison test.

|  | Homeo | Pro-inf | Anti-inf |
| --- | --- | --- | --- |
| Growth rate ($\mu\text{m/s}$) | $0.19 \pm 0.01$ | $0.22 \pm 0.01$ *** | $0.22 \pm 0.01$ ** |
| Shrinkage rate ($\mu\text{m/s}$) | $0.9 \pm 0.01$ | $0.23 \pm 0.01$ ** | $0.22 \pm 0.01$ ** |
| % growth | $15 \pm 1$ | $20 \pm 1$ * | $18 \pm 1$ |
| % shrinkage | $15 \pm 1$ | $16 \pm 1$ | $19 \pm 1$ |
| % pausing | $70 \pm 2$ | $64 \pm 2$ | $63 \pm 2$ |
| Catastrophe frequency ($\text{s}^{-1}$) | $0.16 \pm 0.01$ | $0.13 \pm 0.01$ * | $0.14 \pm 0.01$ |
| Rescue frequency ($\text{s}^{-1}$) | $0.14 \pm 0.06$ | $0.15 \pm 0.01$ | $0.13 \pm 0.01$ |
| MT dynamics ($\mu\text{m/s}$) | $0.06 \pm 0.01$ | $0.08 \pm 0.01$ * | $0.09 \pm 0.01$ ** |
| MTs (n) | 32 | 40 | 36 |
| Cells (n) | 30 | 27 | 29 |

### Homeostatic, pro-inflammatory and anti-inflammatory microglia differ in MT orientation

In most dividing and motile cells, the centrosome is responsible for MT nucleation and anchoring, leading to the formation of radial MT arrays in which all MT minus ends are attached to the centrosome, while MT plus ends extend towards the cell periphery. In contrast, in most differentiated, stationary and axially polarized cells, MTs are more stable and organized in non-centrosomal arrays that are non-radially anchored at the centrosome ^27^.

We hypothesized that in microglia, the transition from the homeostatic phenotype to a migrating reactive state would be paralleled by prominent changes in cell polarity driven by the remodeling of MT anchoring and orientation. To test this, we analyzed the localization and expression of endogenous MT plus end (EB1) and minus end (CAMSAP2) markers in homeostatic, pro-inflammatory and anti-inflammatory microglia cells that had been selected according to their most representative morphology (Figure 2 and S2). While EB1 is a widely adopted marker of actively growing MT plus ends (comets), members of the CAMSAP family regulate the formation and stability of non-centrosomal MT arrays by capping free MT minus ends ^24, 51^. Confocal immunofluorescence analysis showed that in pro-inflammatory and anti-inflammatory microglia, EB1 decorated most free MT ends (Tyr tub stained) (pro-inflammatory, 88%; anti-inflammatory, 82%) that extended towards the cell periphery (Figure 2A), confirming the existence of a prominent pool of dynamic MTs arranged radially with their minus ends attached to a perinuclear region of the cell. EB1 comets were also clearly visible at MT ends in homeostatic microglia although to a lesser extent (homeostatic 63% of MTs p <0.001, see Figure S2A for contingency analysis). Western blot analysis of whole cell lysates showed that total EB1 protein levels did not change in reactive microglia compared to homeostatic cells (Figure S2B and S7). However, a detailed measurement of fluorescence intensity gradients of EB1 positive comets with respect to the location of the cell nucleus (Figure 2B, arrows) identified distinct MT polarity patterns, with anterograde and retrograde orientation. Specifically, a population of retrograde comets was observed in homeostatic and anti-inflammatory microglia as opposed to pro-inflammatory microglia in which all the comets were oriented away from the cell nucleus and toward the cell periphery (23.4%, 12.5% and 0.5%, respectively; p <0.001 Figure S2C). Detection of a pool of retrograde comets in homeostatic microglia suggested the presence of non-centrosomal MT arrays, which we investigated by analyzing the expression and subcellular distribution of endogenous CAMSAP2. Western blot analysis of whole cell lysates revealed endogenous expression of CAMSAP2 in all three microglia phenotypes, with higher protein content in homeostatic microglia (Figure S2D and S7). However, while in homeostatic and anti-inflammatory cells CAMSAP2 often distributed to isolated and clustered puncta along cell ramifications (Figure 2C, and 2D arrows), cytosolic CAMSAP2 signal was detectable only around the perinuclear region in pro-inflammatory microglia (Figure 2C and 2D). Radial profiling of CAMSAP2 fluorescence intensity (Figure 2E), used as a measure of CAMSAP2 distribution in the cytosol, confirmed that CAMSAP2 signal decayed more rapidly at increasing distances from the perinuclear region in pro-inflammatory microglia than in homeostatic and anti-inflammatory cells (Figure 2E, and insert).

**Fig. 2.**
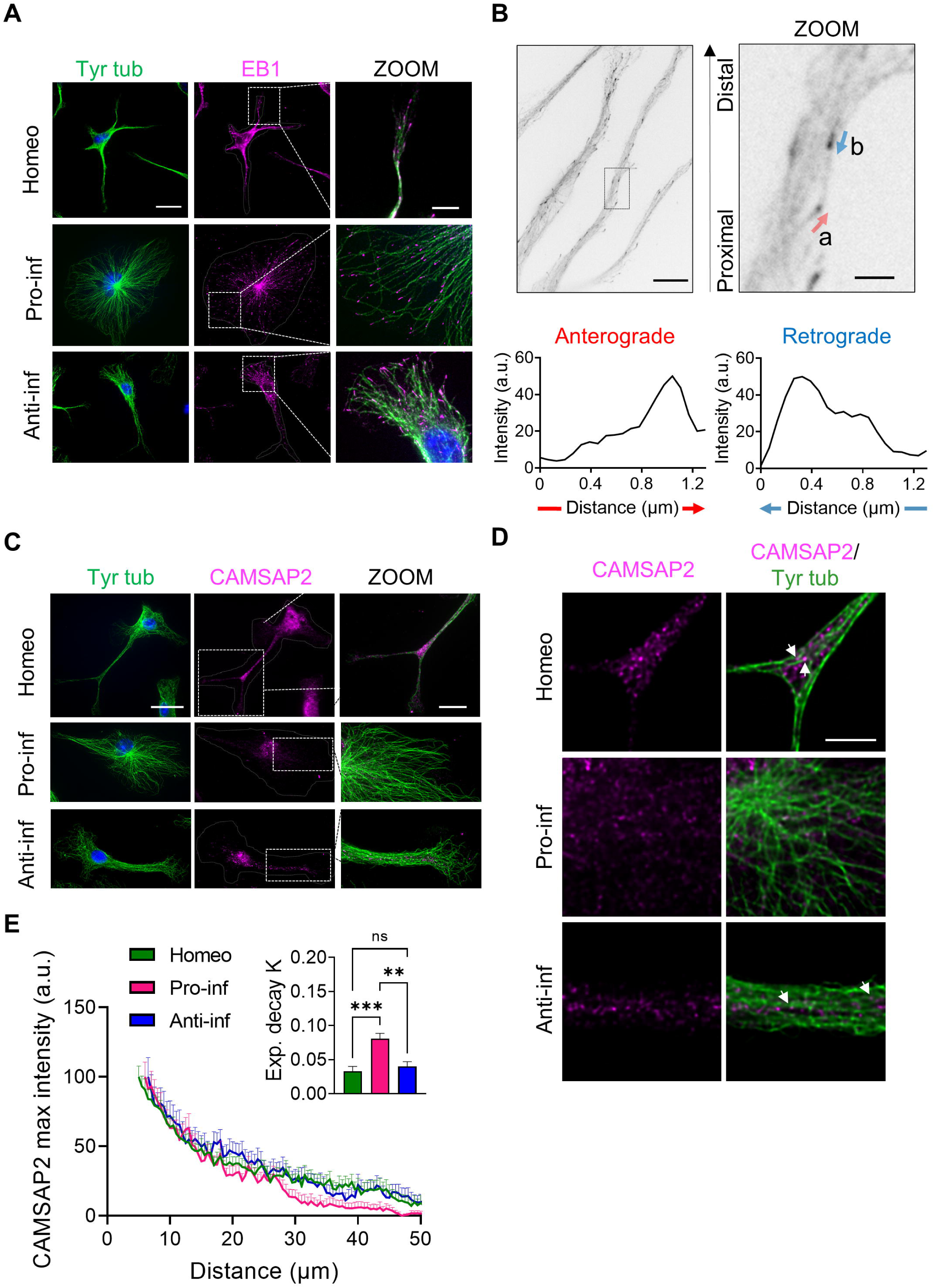
Homeostatic, pro-inflammatory and anti-inflammatory primary microglia differ in MT orientation. **(A)** Representative immunofluorescence images of EB1(magenta) and tyrosinated *α*-tubulin (Tyr tub) (green) in homeostatic (Homeo), pro-inflammatory (Pro-inf) and anti-inflammatory (Anti-inf) microglia. (Scale bar: 20 μm; zoom: 5 μm. Hoechst for nuclei visualization, blue). Cell outlines are indicated by white dashed line. **(B)** Representative inverted contrast single plane image of EB1 immunofluorescence (*top*, *left*. Scale bar 10 μm). Direction of EB1 signal gradient relative to the cell nucleus was used to identify EB1 anterograde (a, red arrow) and retrograde (b, blue arrow) comets (*top*, *right*; scale bar: 2 μm). Bottom: intensity profile of anterograde (left) and retrograde (right) comets. **(C)** Representative z-projection confocal images showing CAMSAP2 (magenta) and Tyr tub (Green) signal in Homeo, Pro-inf and Anti-inf microglia. Cell outlines are indicated by white dashed line (scale bar: 20 μm; zoom: 5 μm. Hoechst for nuclei visualization, blue). **(D)** Single confocal planes at higher magnification of CAMSAP2 (magenta) and Tyr tub (Green) signal in Homeo, Pro-inf and Anti-inf microglia. Scale bar: 5 μm. Note that CAMSAP2 signal is present in microglia processes in Homeo and Anti-inf cells. **(E)** Plot showing maximum fluorescence intensity values of CAMSAP2 vs the radial distance from cell nucleus, obtained with radial profiling, in Homeo (n = 18 cells, green), Pro-inf (n = 14 cells, magenta) and Anti-inf (n = 19 cells, blue) microglia. Curve fit was performed using single exponential decay function. Insert: bar chart reporting the exponential decay constant values (K) for each condition (values are expressed as mean ± SEM from 4 independent experiments; *** p <0.001, ** p <0.01 One-way ANOVA – Tukey’s multiple comparison test.

Altogether, these data indicate that homeostatic and anti-inflammatory microglia display a mixed MT polarity pattern resembling neuronal MTs in dendrites and that the acquisition of a pro-inflammatory phenotype represents a unique example of remodeling of the MT cytoskeleton from an array of parallel non-centrosomal MTs to a radial array of MTs all anchored to pericentrosomal MTOCs through their minus ends.

### Homeostatic microglia nucleate non-centrosomal MTs from Golgi outposts

CAMSAP2 is necessary for the tethering of newly nucleated non-centrosomal MT minus ends during the establishment of polarity in many cell types ^51–53^. We investigated the distribution of γ-tubulin, the major MT nucleator in eukaryotic cells, in homeostatic and reactive microglia by confocal microscopy and found that while in homeostatic and anti-inflammatory cells γ-tubulin displayed a punctate distribution around the perinuclear region and along cellular processes, in pro-inflammatory microglia γ-tubulin signal was restricted to the centrosomal and pericentrosomal area (Figure 3A, Figure S3A and Figure 4 and S4). Radial profiling (Figure 3B) of γ-tubulin fluorescence intensity, used as a measure of γ-tubulin signal distribution in the cell, confirmed that γ-tubulin signal decayed more rapidly at increasing distances from the perinuclear region in pro-inflammatory microglia compared to homeostatic and anti-inflammatory cells (Figure 3B, and insert). This observation was confirmed by the analysis of γ-tubulin signal over the cell area (Figure S3B) and detection of higher γ-tubulin levels in homeostatic and anti-inflammatory microglia compared to pro-inflammatory cells (Figure S3C and S7).

**Fig. 3.**
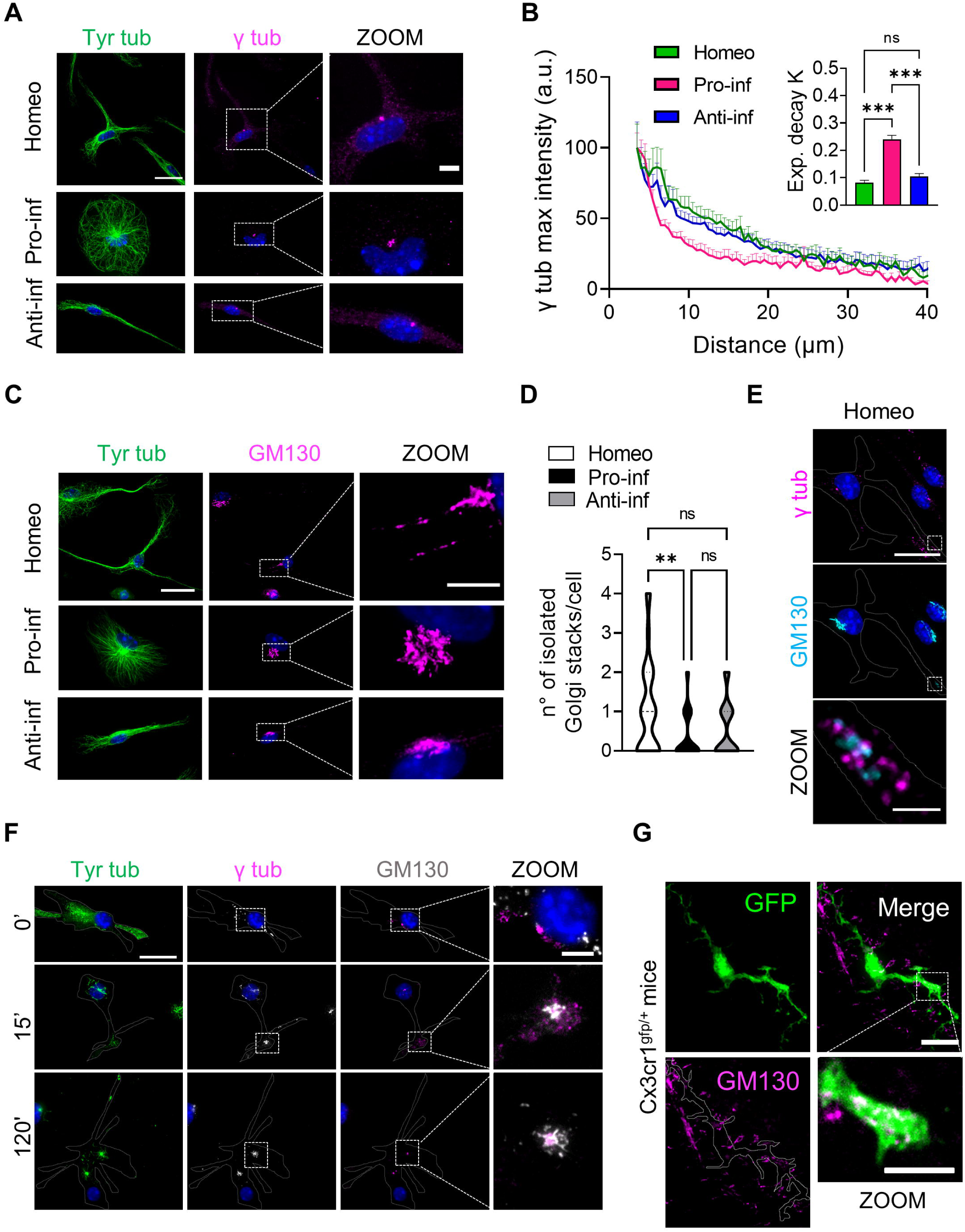
Homeostatic microglia nucleate non-centrosomal MTs from Golgi outposts. **(A)** Representative images of homeostatic (Homeo), pro-inflammatory (Pro-inf) and anti- inflammatory (Anti-inf) microglia stained for tyrosinated *α*-tubulin (Tyr tub, green) and *γ*-tubulin (γ tub, magenta; scale bar: 20 μm; zoom: 5 μm. Hoechst for nuclei visualization, blue). **(B)** Plot showing maximum fluorescence intensity values of γ tub vs the radial distance from cell nucleus, obtained with radial profiling, in Homeo (n = 13 cells, green), Pro-inf (n = 14 cells, magenta) and Anti-inf (n = 14 cells, blue) microglia. Curve fit was performed using single exponential decay function. Insert: bar chart reporting the exponential decay constant values (K) for each condition (values are expressed as mean ± SEM from 3 independent experiments; *** p <0.001, One-way ANOVA – Tukey’s multiple comparison). Note faster decay of γ-tub signal in Pro-inf microglia. **(C)** Representative confocal images showing co-staining of tyrosinated tubulin (Tyr tub) (green) and GM130 (magenta) in Homeo, Pro-inf and Anti-inf microglia (scale bar: 20 μm; zoom: 5 μm. Hoechst for nuclei visualization, blue). **(D)** Violin plot showing number of isolated Golgi stacks per cell in the three phenotypes. Values are expressed as mean ± SEM of Homeo n = 57, Pro-inf n = 34 and Anti-inf n = 36 cells from 3 independent experiments. ** p <0.01. Kuskall-Wallis - Dunn’s multiple comparison test. **(E)** Representative images showing GM130 (cyan) and γ-tub (magenta) staining in Homeo microglia. (Scale bar: 20 µm; zoom: 2 µm. Hoechst for nuclei visualization, blue). Note the presence of Golgi outposts in microglia processes. **(F)** Representative confocal images of the time course of the MT re-nucleation assay after nocodazole washout in Homeo microglia stained for Tyr tub (green), GM130 (gray) and γ tub (magenta). Scale bar: 20 µm; zoom: 5 µm. Hoechst for nuclei visualization, blue). Time 0’ represents the MT depolymerizing effect of nocodazole in homeostatic cells without free tubulin extraction. Note that MTs nucleate from distal Golgi outposts that are positive for *γ*-tubulin. **(G)** Representative z- projection confocal images of retinal slices (50 μm thickness) from cx3cr1^gfp/+^ mice, expressing GFP in microglia cells, stained with GM130 (magenta) to visualize Golgi outposts. Scale bar: 5 μm. Zoom is a single confocal plane of a microglia ramification stained for GM130; scale bar: 5 μm.

**Fig 4.**
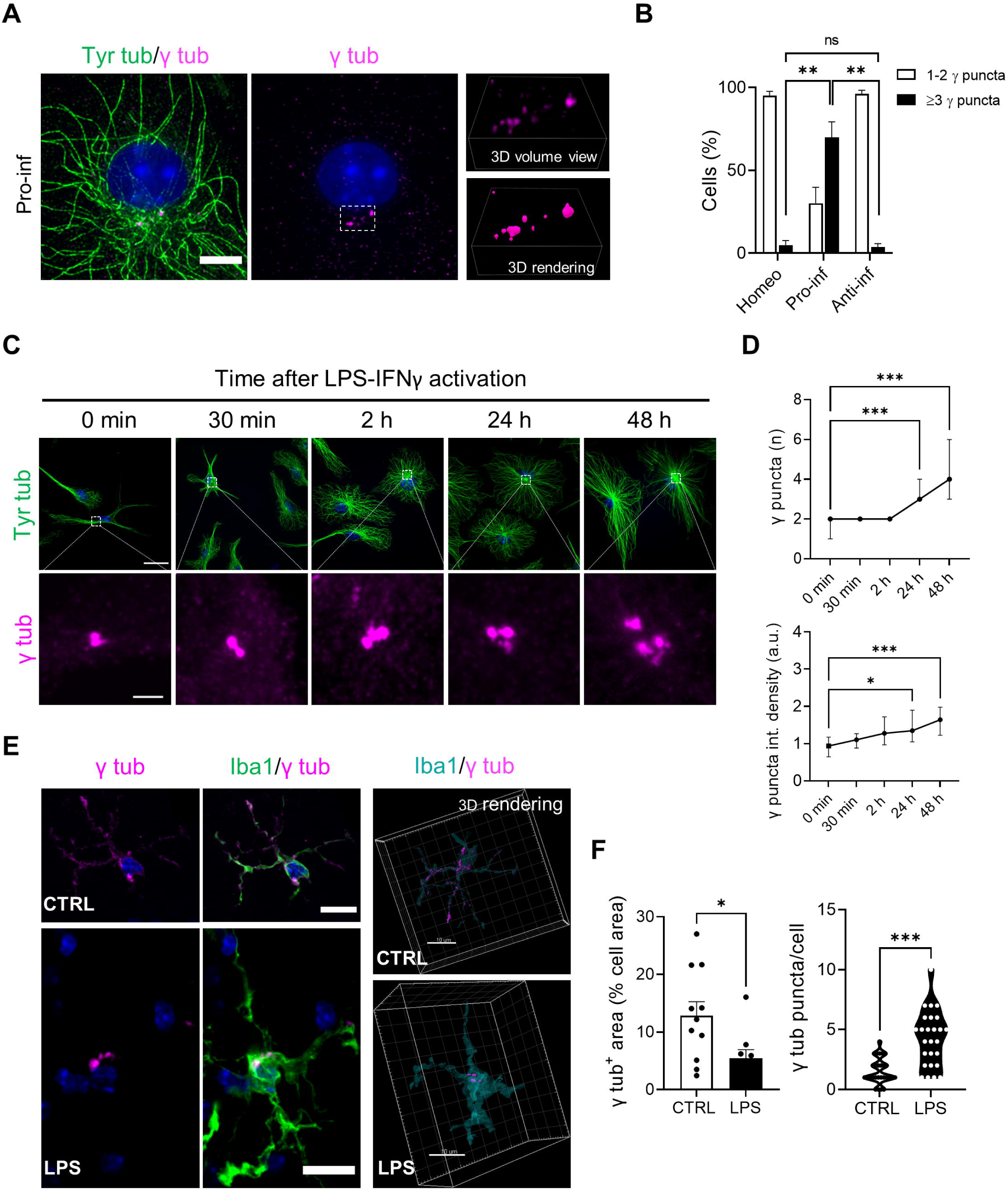
Redistribution of pericentriolar material is a hallmark of pro-inflammatory microglia *in vitro* and *in vivo.* **(A)** Representative confocal images showing *γ*-tubulin (γ tub) puncta (magenta) and tyrosinated *α*-tubulin (Tyr Tub, green) immunolabeling in pro-inflammatory (Pro- inf) microglia (*left*, *middle*). Scale bar: 5 µm. Hoechst for nuclei visualization, blue. *Right:* relative volume view *(top)* and 3D rendering *(bottom)* of γ tub puncta (magenta) acquired via structured illumination microscopy. **(B)** Bar chart reporting the percentage of cells displaying 1-2 γ tub puncta (white bars) or >3 γ tub puncta (black bars) in homestatic (Homeo), pro-inflammatory (Pro-inf) and anti-inflammatory (Anti-inf) microglia. Values are expressed as mean ± SEM from 3 independent experiments. ** p <0.01. One-way ANOVA - Dunnett’s multiple comparison test. **(C)** Time course of γ -tubulin (γ tub) redistribution during the process of microglia pro- inflammatory activation: representative images showing γ tub (magenta) and tyrosinated *α* - tubulin (Tyr tub, green) staining at different time points (0 min, 30 min, 2 h, 24 h, 48 h). Scale bar: 20 µm; zoom, 2 µm. Hoechst for nuclei visualization, blue. **(D)** Time course of the number of γ puncta per cell (top) and the quantification of γ puncta fluorescence intensity (bottom) during the process of microglia pro-inflammatory activation. Values are expressed as median ± interquartile range (T0 n = 32; T30 min n = 40; T2h n = 44; T24h n = 41; T48h n = 33; from 3 independent experiments); *** p <0.001; ** p <0.01; * p <0.05, Kruskal-Wallis test - Dunn’s multiple comparison test respect to T0. **(E)** *Right*: representative maximum intensity projections of retinal slices (50 μm thickness) from control (CTRL) mice stained for γ tub (magenta) and Iba-1 (green) antibodies (Scale bar: 10 µm. Hoechst for nuclei visualization, blue). *Left*: 3D rendering of same retinal microglia highlighting the intracellular distribution of γ tub in CTRL (sham) and LPS treated mice. Scale bar: 10 μm. **(F)** *Left*: scatter dot plot showing the γ tub signal over the cell area of retinal microglia from CTRL (sham) and LPS treated mice. Values are expressed as mean ± SEM of n = 11/3 cells/mice (CTRL) and n = 9/3 cells/mice (LPS). * p <0.05. Student’s t-test. *Right*: violin plot showing the number of γ tub puncta per microglia in retinal slices from CTRL and LPS treated mice of n = 28/3 cells/mice (CTRL) and n = 26/3 cells/mice (LPS). *** p <0.001, Student’s t-test.

Altogether, these data demonstrate that the acquisition of a pro-inflammatory phenotype is characterized by restricted localization of γ-tubulin to a pericentrosomal area, which is necessary to establish radial MT arrays. The presence of γ-tubulin in cell ramifications of homeostatic microglia further suggests that microglia are alternatively enriched in non-centrosomal MT nucleation sites, which are necessary to establish non-centrosomal MT arrays.

Golgi outposts can serve as acentrosomal MTOCs in other highly polarized brain cells, such as neurons and oligodendrocytes ^54–56^. We thus analyzed the distribution of the Golgi marker GM130, a scaffolding protein peripherally associated with Golgi membranes and a marker of Golgi outposts ^57^. Co-staining of GM130 with tyrosinated tubulin (Tyr tub) demonstrated that the presence of Golgi outposts is a feature of homeostatic microglia, and that their presence was dramatically reduced in pro-inflammatory cells (Figure 3C and 3D). Importantly, most of the isolated GM130 positive mini-stacks (73.6%) were decorated by γ-tubulin (Figure 3E). To determine whether Golgi outposts could function as MTOCs in homeostatic microglia, MT nucleation was evaluated *in situ* by analyzing MT re-nucleation after nocodazole washout (Figure S3D). We found that both at 15 and 120 mins after nocodazole washout to allow MT-regrowth after nocodazole-induced MT depolymerization, MTs (stained with Tyr tub) emerged from Golgi membranes (GM130 positive) both at pericentrosomal sites and at Golgi outposts located far from the centrosome (Figure 3F). Importantly, *γ*-tubulin was localized to nucleation-competent Golgi outposts, indicating that non-centrosomal MT re-nucleation did not occur spontaneously at these sites but was strictly dependent on the presence of a *γ*-tubulin nucleation complex (Figure 3E).

These data demonstrate that in homeostatic microglia Golgi outposts can function as sites of acentrosomal MT nucleation and suggest that *γ*-tubulin-dependent non-centrosomal nucleation is necessary to establish an asymmetric MT array in these cells.

To assess whether these *in vitro* observations were representative of MT nucleation in microglia residing in tissue, we analyzed the subcellular distribution of GM130 in retinal microglia: retina and brain share a common embryological origin and similar cell types ^58–63^. Moreover, retinal neurons are arranged in distinct layers, and microglia are usually restricted to the retinal ganglion cell layer, thus offering an accessible structure for the imaging of their MT cytoskeleton. To identify the microglial cytoskeleton in retina we used cx3cr1^gfp/+^ mice, which constitutively express GFP in microglia. As expected, retinal microglia from control cx3cr1^gfp/+^ mice displayed a highly ramified morphology (Figure S3E), typical of homeostatic surveillant cells ^6, 64, 65^. More importantly, confocal immunofluorescence analysis of GM130 signal in retinal GFP positive microglia confirmed the presence of isolated Golgi outposts also in the processes of homeostatic microglia residing in tissue (5 ± 1 per cell, n = 11; Figure 3G, S3F and S3G).

Altogether, these results support the notion that MT organization in homeostatic microglia resembles the MT architecture typical of highly polarized, terminally differentiated cells and strongly suggest that γ-tubulin dependent non-centrosomal MT nucleation at Golgi outposts is a *bona fide* feature of homeostatic, surveilling microglia *in vitro* and *in vivo*.

### Pericentrosomal redistribution of microtubule-nucleating material is a hallmark of pro-inflammatory microglia and regulates IL-1β secretion

The recruitment of pericentriolar material (PCM) to the centrosome has been described as a functional step for macrophage activation upon pro-inflammatory stimuli ^66^. We thus investigated whether the recruitment of γ-tubulin to a pericentrosomal area was also a hallmark of pro-inflammatory microglia.

As revealed by super-resolution microscopy, pro-inflammatory microglia exhibited multiple γ-tubulin^+^ puncta that localized to a perinuclear region (Figure 4A). Quantification of the number of γ-tubulin^+^ puncta indicated that most pro-inflammatory cells had more than 3 puncta (70 ± 10%; Figure 4B). Conversely, almost all homeostatic and anti-inflammatory microglia displayed only 1 or 2 γ-tubulin^+^ puncta (95 ± 3% and 96 ± 2%, respectively; Figure 4B). γ-tubulin localization to pericentrosomal puncta showed a time-dependent increase of both number and fluorescence integrated density upon LPS-INFγ challenge (Figure 4C and 4D) and the recruitment of γ-tubulin^+^ puncta was dependent on a dynamic MT cytoskeleton because a low dose of taxol was sufficient to inhibit it (Figure S4A and S4B). Notably, in most pro-inflammatory cells with >2 *γ*-tubulin^+^ puncta (65 ± 10%; Figure S4C) the centrosomal marker centrin-3 localized only to ≤2 *γ*-tubulin^+^ puncta. In addition, while a quarter of pro-inflammatory cells was proliferating (23 ± 4%; Figure S4E and S4F), most pro-inflammatory microglia displayed >2 γ-tubulin^+^ puncta (70 ± 10%; Figure 4B) and ≤2 centrin^+^ puncta (Figure S4D). Conversely, PCM localization to *γ*- tubulin^+^ puncta was confirmed by coimmunostaining with pericentrin (86 ± 5% of colocalizing puncta), a conserved PCM scaffold protein necessary for MTOC assembly and maturation ^67^ Figure S4G). Indeed, both centrin^+^ and centrin^-^ *γ*-tubulin^+^ puncta localized to the center of MT asters (Fig. S4H) and MT re-growth after nocodazole washout (Figure S4I,J) revealed that *de novo* MT nucleation occurred at *γ*-tubulin^+^ puncta (Figure S4L). This indicated that *γ*-tubulin reorganization in pro-inflammatory microglia is ascribed to PCM maturation that is uncoupled from cell or centrosome duplication, and that *de novo* generated PCM puncta act as MTOCs.

Perinuclear γ-tubulin redistribution was also observed in proinflammatory microglia in a mouse model of retinal inflammation. For this, we took advantage of a well-established protocol of acute inflammatory uveitis ^32–37^ induced by intravitreal injection of LPS (Figure S5A) to activate retinal microglia towards the pro-inflammatory phenotype. Microglia residing in retinal slices from LPS-treated mice acquired an amoeboid morphology with reduced branching complexity, as revealed by skeleton analysis of Iba1 positive cells (Figure S5B and S5C). Moreover, co-immunolabelling with Iba1 and γ-tubulin demonstrated that while in control (sham) mice retinal microglia displayed punctate diffuse γ-tubulin staining along cellular ramifications (Figure 4E), in LPS treated mice microglia clearly exhibited a condensed γ-tubulin pattern, clustered around a perinuclear region (Figure 4E). This was confirmed by quantitative analysis of γ-tubulin signal over the cell area and of the number of γ-tubulin^+^ puncta per cell (Figure 4F).

In summary, these data demonstrate that the presence of pericentrosomal MTOC maturation is a *bona fide* feature of pro-inflammatory microglia *in vitro* and *in vivo*.

Next, we examined if maturation of pericentrosomal MTOCs was a regulatory step for microglia acquisition of the pro-inflammatory phenotype. To this end, we treated microglia with the selective PLK4 inhibitor centrinone to hamper PCM maturation ^68^ prior to LPS-INFγ stimulation (Figure 5A) and measured its effects on EV blebbing and cytokine release (Figure 5). First, we found that centrinone increased the number (4.7-fold; Figure 5C, top) and reduced the diameter (by 30%; Figure 5C, bottom) of EVs blebbing from the cell surface of pro-inflammatory microglia (Fig. 5A-C). Activation of NLRP3-inflammasome is a characteristic feature of pro-inflammatory microglia and macrophages ^69–73^ and the centrosome-associated protein kinase NEK7 is required for NLRP3–inflammasome activation (for review see ^62^). In addition, NLRP3 activation is required for IL-1*β* maturation in immortalized and primary microglia ^69, 75^, and inhibition of PLK4 leads to NLRP3 hyper-activation through NEK7 dephosphorylation in bone marrow-derived macrophages ^68^. We then measured NLRP3-inflammasome protein levels in homeostatic and pro-inflammatory microglia upon centrinone treatment. We found that in homeostatic microglia, LPS-IFNγ but not centrinone alone increased NLRP3 expression while centrinone / LPS-IFNγ co-treatment enhanced NLRP3 more than LPS-IFNγ alone (Figure 5D and S7), suggesting that inhibition of PCM maturation induces NLRP3 hyperactivation in pro-inflammatory microglia. We investigated whether PLK4 inhibition modulates microglia IL- 1*β* release and found that co-treatment with centrinone enhanced pro-inflammatory microglia-mediated IL-1*β* release, without affecting IL-1*β* release from homeostatic cells (Figure 5E). In addition, we observed that centrinone did not modulate the release of IL-10 and IL-6 (Figure 5F), indicating that PLK4 inhibition enhances pro-inflammatory microglia IL-1*β* release through NLRP3-inflammasome activation but has no effect on NLRP3-independent pathways.

**Fig 5.**
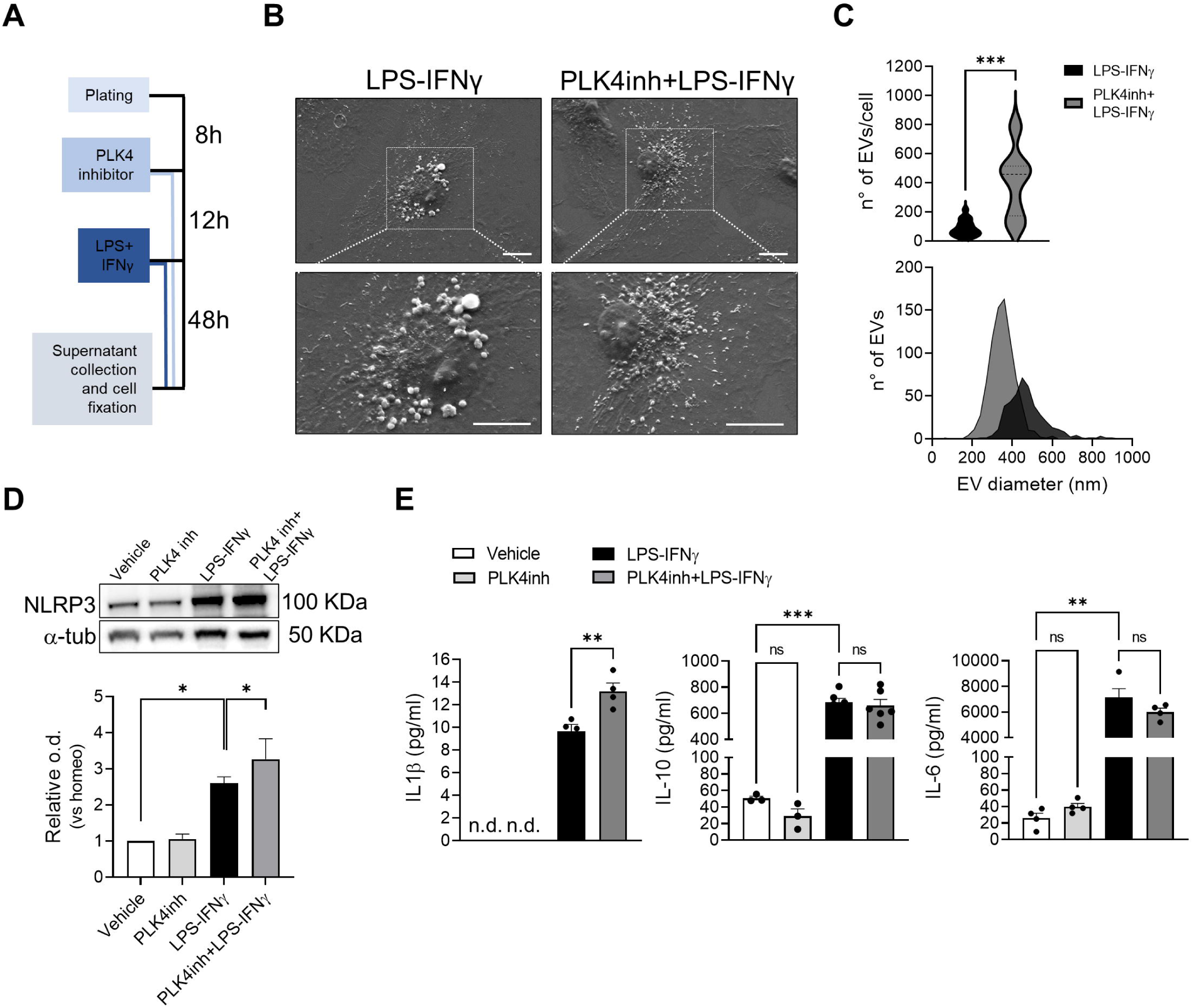
Inhibition of pericentriolar material maturation induces NLRP3 inflammasome hyperactivation in pro-inflammatory microglia. (**A**) Experimental timeline of PLK4 inhibitor treatment and pro-inflammatory cytokine administration. (**B**) Scanning electron micrographs showing extracellular vesicles (EVs) on the surface of LPS-IFNγ (*left*) and PLK4 inhibitor+LPS-IFNγ (*right*). Scale bar: 10 μm. (**C**) *Top*: violin plot showing the number of EVs in Pro-inf microglia with or without PLK4inh treatment (LPS-IFNγ n = 17 cells, PLK4 inhibitor+LPS-IFNγ n = 19 cells from 2 independent cultures; *** p <0.001, Student’s t-test). *Bottom*: distribution of EV size measured on cell surface of LPS-IFNγ and PLK4 inhibitor+LPS-IFNγ microglia (LPS-IFNγ n = 17 cells, PLK4 inhibitor+LPS-IFNγ n = 19 cells). (**D**) *Bottom*: bar chart reporting the amount of NLRP3 protein level in Vehicle, PLK4 inhibitor, LPS-IFNγ and PLK4 inhibitor+LPS-IFNγ microglia; *top*: representative immunoblot of NLRP3. Values are expressed as median ± interquartile range from 4 independent experiments, * p <0.05, Mann Whitney test. (**E**) Scatter dot plot showing protein quantification by ELISA of *left*: IL1*β* (n = 4 independent experiments, n.d. = non detectable), *middle*: IL-10 (Vehicle n = 3, PLK4inh n = 3, LPS-IFNγ n = 5 and PLK4 inhibitor+LPS-IFNγ n = 5 independent experiments) and *right*: IL-6 (n = 4 independent experiments). Values are expressed as mean ± SEM. For IL- 1*β*: ** p <0.01, Student’s t-test. Note that supernatants from both Vehicle and PLK4 inhibitor microglia have undetectable levels of cytokine. For IL-10 and IL-6: *** p <0.001, ** p <0.01; One-way ANOVA - Holm-Šídák’s multiple comparison test.

Together, these data demonstrate that microglia pro-inflammatory reactivity induces PCM maturation *in vivo* and *in vitro*, and that this step is a negative regulator of NLRP3-dependent IL- 1*β* release. They also indicate that remodeling of the MT cytoskeleton during pro-inflammatory reactivity is temporally coupled to cytokine release, providing a novel potential target of microglia treatment in inflammatory diseases.

## DISCUSSION

Here we describe the reorganization of the microglial MT cytoskeleton that characterizes the transitions between homeostatic, pro-inflammatory and anti-inflammatory states, and demonstrate the functional interplay between microglial PCM maturation and pro-inflammatory reactivity. Our findings demonstrate that pro-inflammatory microglia reactivity orchestrates a so far unique rearrangement of the MT cytoskeleton from a non-centrosomal array of parallel and stable MTs nucleated at Golgi outposts characteristic of the homeostatic state, to a radial array in which MTs are anchored to *de novo* formed pericentrosomal MTOCs through their minus ends. Through *in vitro* phenotyping and *in vivo* validation, we report four main findings summarized in Figure S6: 1) Homeostatic microglia possess stable MT arrays, while microglia reactivity increases MT dynamic behavior. 2) Non-centrosomal MT organization in arrays with mixed polarity is a feature of homeostatic microglia, like the architecture typical of highly specialized cells such as neurons and oligodendrocytes. 3) Pro-inflammatory microglia reactivity results in restricted *γ*- tubulin localization to puncta around the centrosome because of *de novo* PCM and MTOC maturation, providing a novel distinct marker of microglia reactivity in live-imaging studies 4) PCM maturation in pro-inflammatory microglia is a regulator of NLRP3-dependent IL-1*β* release. To date, only circumscribed evidence has suggested that ramified microglia possess more acetylated and detyrosinated MTs than ameboid pro-inflammatory cells ^14, 76^. Here, we show that homeostatic ramified microglia display higher levels of tubulin acetylation and detyrosination, two indirect indicators of “older”, i.e., more stable and less dynamic, MT subpopulations ^49, 77^. Moreover, we report that during classical and alternative activation obtained with either LPS-IFNγ or IL-4 stimulation respectively ^78–80^, microglia MTs become less stable and more dynamic, suggesting that the acquisition of new cellular functions induces changes in MT stability *via* modulation of MT dynamics ^81^.

We describe that homeostatic/ramified microglial MTs exhibit an asymmetric dendrite-like organization characterized by EB1 comets arranged in mixed polarity with the minus-end capping protein CAMPSAP2 localized at branching points and cell ramifications. In anti-inflammatory microglia, we find lower levels of CAMSAP2 that localize in a similar fashion in bipolar processes. The presence of CAMSAP2 in microglia processes might suggest that, as in neurons ^51, 82–86^, stabilization of non-centrosomal MTs at their minus ends is important to achieve elongated bipolar morphology, typical of anti-inflammatory microglia, and for the formation of long branched cellular extensions patrolling brain parenchyma in homeostatic microglia.

We find that the acquisition of a pro-inflammatory phenotype disrupts the cellular asymmetry of homeostatic microglia and reduces the pool of non-centrosomal, parallel and mixed oriented MTs, leading to their rearrangement into a radial array of uniformly oriented MTs characteristic of the ameboid shape. In addition, while in pro-inflammatory microglia all the MTs are anchored to a centrosomal region, homeostatic microglia nucleate acentrosomal MTs from Golgi outposts located far from the cell body at the branching points of microglia ramifications, resembling the structure of the dendritic tree of mature neurons ^51, 87, 88^ or the organization of oligodendrocytic myelin sheaths ^55, 56^. This Golgi outpost-dependent non-centrosomal nucleation contrasts to pro-inflammatory and anti-inflammatory microglia in which the Golgi apparatus displays a compact perinuclear location ^89^, suggesting that in reactive microglia cellular arborization is reduced by restricting the Golgi to a region adjacent to the centrosome, which acts as the major MT nucleator in these cells. Indeed, we observed that the acquisition of a pro-inflammatory phenotype is characterized by γ-tubulin redistribution to puncta located to a pericentrosomal region, a feature we confirmed in retinal microglia residing in tissue. Co- localization of *γ*-tubulin with pericentrin and *de novo* nucleation of radial MTs from *γ*-tubulin^+^ puncta upon nocodazole washout strongly suggests that these protein assemblies are composed of PCM and act as pericentrosomal MTOCs. Importantly, γ-tubulin redistribution during the transition to a pro-inflammatory phenotype did not derive from centriolar duplication during cell division ^90, 91^ as no more than 2 γ-tubulin^+^ puncta colocalized with centrin-3 in the same cell, an abundant protein associated with the centrosome ^92^. In addition, γ-tubulin redistribution was strictly dependent on a dynamic MT cytoskeleton, suggesting that the increase in MT dynamicity is necessary for the relocation of MT nucleating material to the centrosome. Importantly, our observations indicate that pericentrosomal redistribution of microtubule nucleating material may further provide a distinct and highly valuable live-imaging marker of microglia activation to detect progression of neuroinflammatory disease and efficacy of therapeutics over time.

We find that PCM maturation occurring in pro-inflammatory microglia negatively regulates IL-1*β* release through a non-classical secretory pathway ^93^. Indeed, inhibition of PLK4 during the acquisition of a pro-inflammatory phenotype reduces pericentrosomal γ-tubulin^+^ puncta formation, increases NLRP3 expression, and potentiates IL-1*β* release without affecting the release of IL-6 and IL-10. NLRP3-inflammasome hyperactivation is consistent with NLRP3 hyperactivation through NEK7 dephosphorylation by inhibition of PLK4 in macrophages ^68, 74^. Given that the NLRP3-inflammasome mediates interleukin activation ^94^, NLRP3-inflammasome hyperactivation may account for the increase in IL-1*β* release and the accumulation of blebbing microvesicles we observe in microglia stimulated *in vitro*. Interestingly, our results on cytokine release upon PLK4-inhibition differ from those reported after long-term (7 days) inhibition in macrophages activated with LPS ^66^ suggesting that the response to loss of PLK4 activity is cell type specific or longer treatments may have either secondary or opposite effects. Further studies are necessary to discriminate between these possibilities.

In summary, we identify a heretofore unique example of MT reorganization from a non-centrosomal array of MTs with mixed polarity to a radial array in which all the MTs are uniformly oriented and anchored either at the centrosome or pericentrosomal MTOCs. Our structural, functional and in tissue analyses further demonstrate that acentrosomal MT nucleation at Golgi outposts may play an important role in supporting the patrolling phenotype of microglia cells, that tubulin remodeling enables microglia reactivity *in vitro* and in tissue, and that targeting PCM maturation in reactive microglia may represent a new approach to limit tissue damage during neurodegenerative disease in which microgliosis contributes to neuronal injury and cognitive decline ^2, 12^. In addition, given the newly identified role for a population of spinal CD11c^+^ microglia in the remission and recurrence of neuropathic pain ^95^, it will be become critical to determine the contribution of spinal microglial MT dysfunction in the peripheral neuropathy caused by chemotherapeutic drugs, most of which target the MT cytoskeleton.

## Acknowledgments

The authors wish to thank the Animal Facility of Physiology and Pharmacology Department of Sapienza University. The graphical abstract in Figure S6 was created using BioRender.com. We are grateful to David Sulzer for comments on the manuscript and useful discussions. Further information and requests for resources and reagents should be directed to and will be fulfilled by the lead contact, Silvia Di Angelantonio.

This study was supported by NIH/NIA R56AG050658 and NIH/NINDS R21 NS120076-01 grants to FB and a Fulbright Award to FB and SDA. This research was also funded by the CrestOptics- IIT JointLab for Advanced Microscopy (to SDA and GCR); the Regione Lazio MARBEL Life2020 and Bio3DBrain FSE 2014-2020 grants (to SDA); by Sapienza University RM118163E0297F84 and PH12017270934C3C grants (to SDA) and Fondazione Istituto Italiano di Tecnologia (to MR and CS). CS was also supported by the Ph.D. program in Life Science at Sapienza University in Rome. The research leading to these results has been also supported by European Research Council Synergy grant ASTRA (n. 855923 to GCR.).

## Author contributions

FB, SDA, and MR designed the study and wrote the manuscript. CS, MR, MM, DD, FC, GP, MG and AI performed experiments and generated primary data. GG generated code; CS, MR, FC, DD, GP, MM, MG and SDA performed data analysis. EDL, SDP, GCR, DR, and RG provided helpful insights and contributed key techniques. FB and SDA supervised data analysis and experiments.

## Data and materials availability

This study did not generate new unique reagents. The data that support the findings of this study are available from the corresponding author upon reasonable request. The code is available at https://github.com/ggosti/maxIntensityRadialProfile.

