## Supplementary Figures & Legends for "Microglia reactivity entails microtubule remodeling from acentrosomal to centrosomal arrays"

Fig.S1

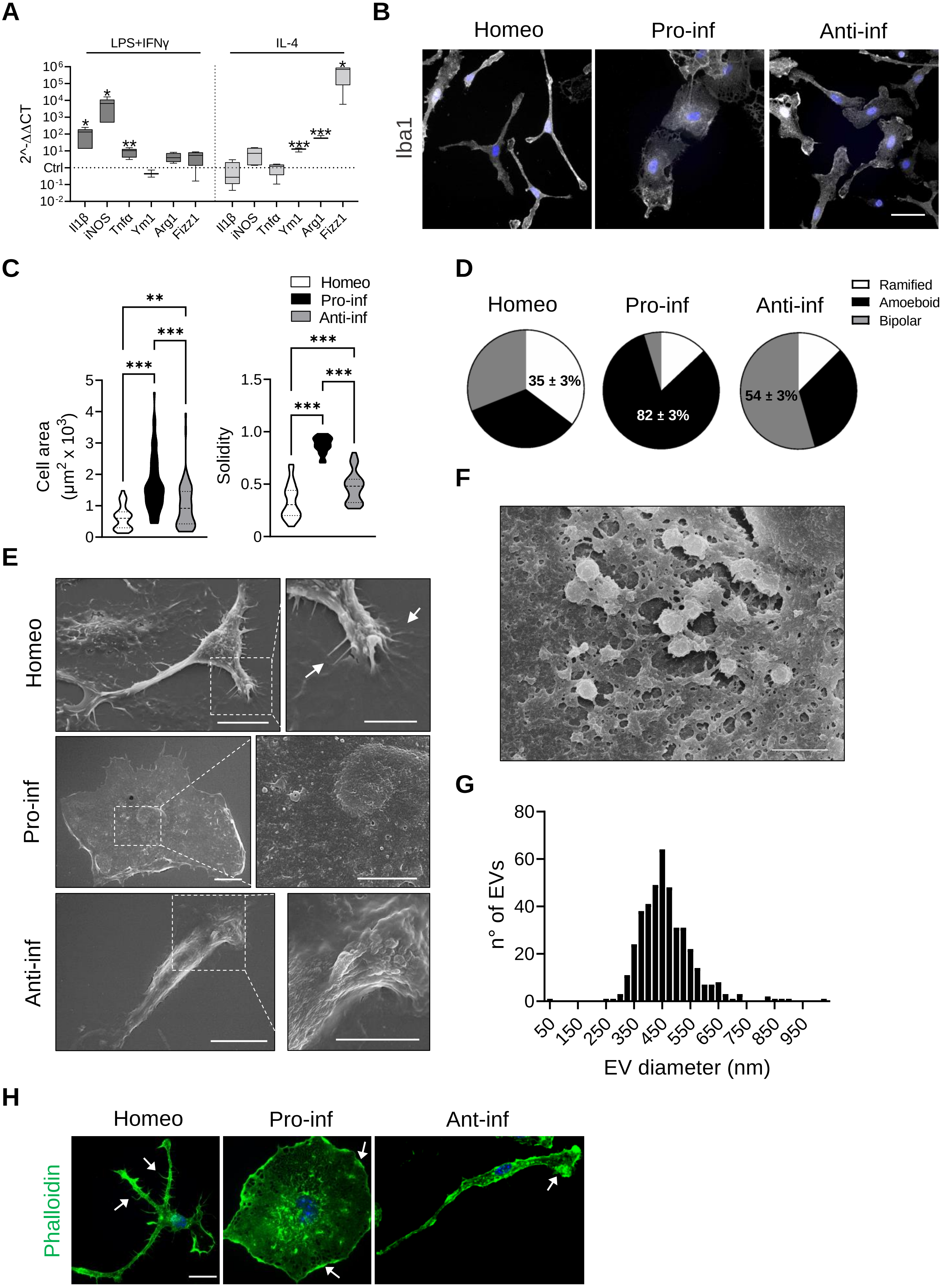

A
LPS+IFNγ
IL-4
2^-ΔΔCT
Ctrl
Il1β
iNOS
Tnfα
Ym1
Arg1
Fizz1
B
Homeo
Pro-inf
Anti-inf
Iba1
C
Homeo
Pro-inf
Anti-inf
Cell area (μm² x 10³)
Solidity
D
Ramified
Amoeboid
Bipolar
35 ± 3%
82 ± 3%
54 ± 3%
E
F
G
n° of EVs
EV diameter (nm)
H
Ant-inf
Phalloidin

**Fig. S1. Molecular and morphological characterization of homeostatic, pro-inflammatory and anti-inflammatory primary microglia**

**(A)** RT-qPCR reveals increased expression of pro-inflammatory (*Il1b, iNOS, Tnfα*) and anti-inflammatory genes (*Ym1, Arg1, Fizz1*) upon LPS-IFNγ or IL-4 challenge, respectively. Gene expression is normalized to the housekeeping gene *Gapdh,* n = 4 independent cultures. Box plots indicate the median, 25th, and 75th percentiles; whiskers show the minimum and maximum. *** $p < 0.001$; ** $p < 0.01$; *$p < 0.05$, Student's t-test. **(B)** Representative images showing Iba1 immunostaining (gray) of microglia in CTRL (Homeo) condition and following LPS-IFNγ (Pro-inf) or IL4 (Anti-inf) treatment (Scale bar: 20 µm. Hoechst for nuclei visualization, blue). **(C)** Violin plots showing cell surface area (*left*) and solidity coefficient (*right*) related to Homeo (white), Pro-inf (black) and Anti-inf (gray) cells (n = 70 cells for each condition, from 3 independent experiments. *** $p < 0.001$; **$p < 0.01$. Kruskall-Wallis - Dunn's multiple comparisons test for cell area, *left*; One-way ANOVA - Tukey's multiple comparison test for solidity, *right*). **(D)** Pie charts illustrating the distribution of cell morphology in Homeo, Pro-inf and Anti-inf conditions. Ramified cells are enriched in Homeo (35 ± 3%), amoeboid in Pro-inf (82 ± 3%) and bipolar in Anti-inf (54 ± 3%), n = 3 independent experiments. **(E)** Representative scanning electron micrographs of microglia cells in Homeo (*top*), Pro-inf (*middle*) and Anti-inf (*bottom*) conditions. Ramified cells (Homeo) show filopodia extensions as indicated by arrows, ameboid cells (Pro-inf) exhibit numerous extracellular vesicles on the cell surface, while bipolar cells (Anti-inf) are characterized by extensive membrane ruffling on the cell surface and leading edge. Scale bar: 10 µm; zoom, 5 µm. **(F)** Scanning electron micrographs showing extracellular vesicles on the Pro-inf cell surface at higher magnification. Scale bar: 2 µm. **(G)** Distribution of EVs size measured on ameboid cell surface. n = 22 cells. **(H)** Representative confocal images of microglia labelled with Alexa Fluor 488-conjugated phalloidin (green). Ramified cells (Homeo) exhibit filopodia like structures (as indicated by arrows) while ameboid (Pro-inf) and bipolar cells (Anti-inf) are characterized by membrane ruffles along cell borders and at lamellipodia (as indicated by arrows). Scale bar: 25 µm. Hoechst for nuclei visualization, blue.

**Fig.S2**

**A**

| | Tyr+/EB+ comets | Tyr+/EB- comets | TOT |
|---|---|---|---|
| Homeo | 96 | 57 | 153 |
| Pro-inf | 361 | 48 | 409 |
| Anti-inf | 146 | 33 | 179 |
| TOT | 603 | 138 | 741 |

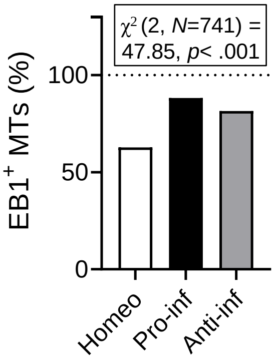


**B**

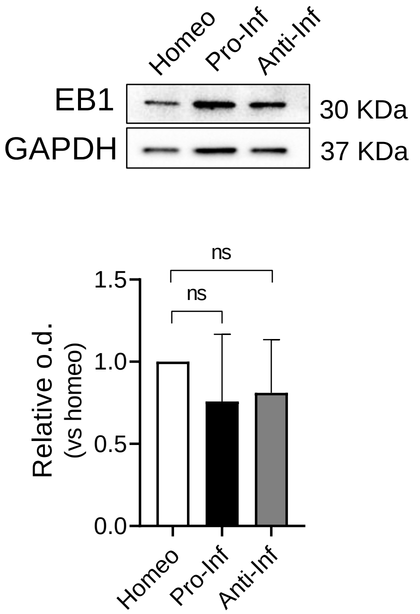


**C**

| | Retrograde comets | Anterograde comets | TOT |
|---|---|---|---|
| Homeo | 33 | 108 | 141 |
| Pro-inf | 1 | 187 | 188 |
| Anti-inf | 32 | 224 | 256 |
| TOT | 66 | 519 | 585 |

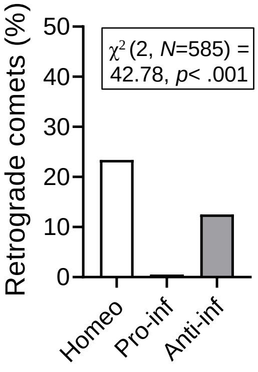


**D**

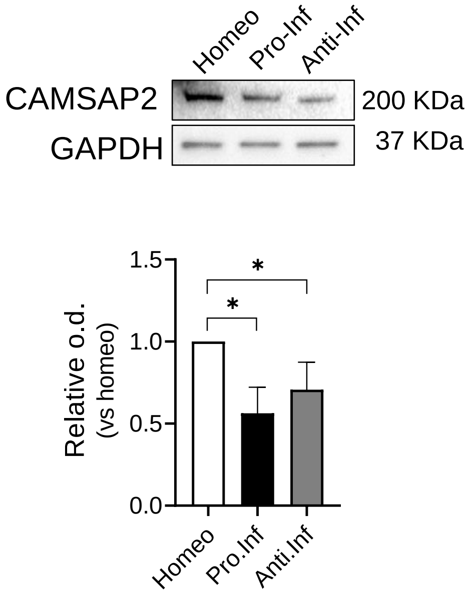

**Fig. S2. Analysis of EB1 and CAMSAP2 expression in homeostatic, pro-inflammatory and anti-inflammatory microglia**

**(A)** Contingency analysis of EB1 and tyrosinated a-tubulin (Tyr tub) co-staining in homeostatic (Homeo), pro-inflammatory (Pro-inf) and anti-inflammatory (Anti-inf) microglia: percentage of double positive staining are reported in the bar chart, $\chi^2$ parameters are reported in the insert. (**B**) *Bottom*: bar chart reporting the amount of EB1 protein level in microglia phenotypes; *top*: representative immunoblot of EB1. Values are expressed as median ± interquartile range from 4 independent experiments. p=0.31, Mann Whitney test. (**C**) Contingency analysis of comets in Homeo, Pro-inf and Anti-inf microglia: percentage of retrograde comets are reported in the bar chart, $\chi^2$ parameters are reported in the insert. (**D**) *Bottom*: bar chart reporting the amount of CAMSAP2 protein level in microglia phenotypes; *top*: representative immunoblot of CAMSAP2. Values are expressed as median ± interquartile range from 4 independent experiments. * $p < 0.05$, Mann Whitney test.

Fig.S3

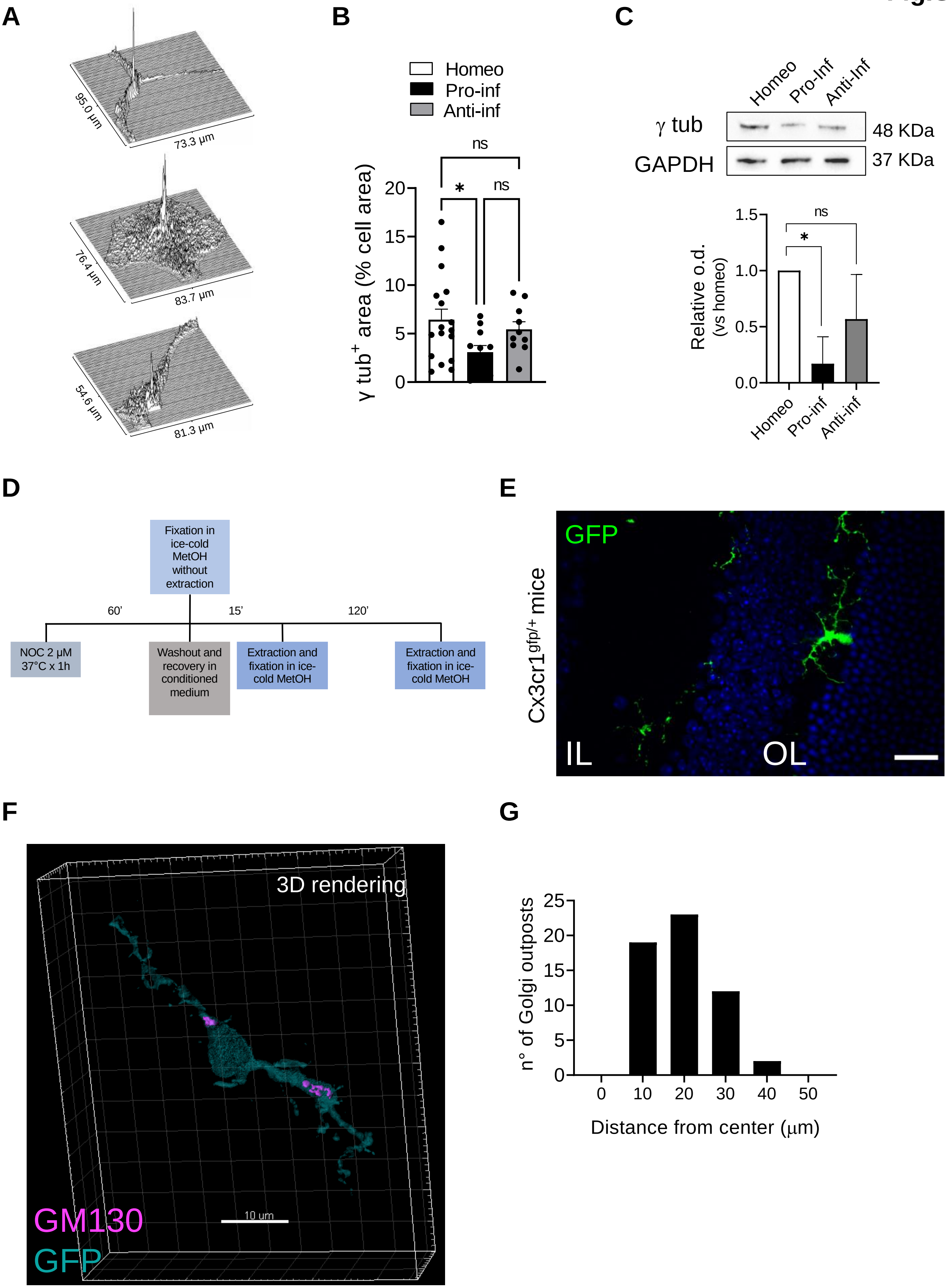

A
95.0 µm
73.3 µm
76.4 µm
83.7 µm
54.6 µm
81.3 µm
B
Homeo
Pro-inf
Anti-inf
ns
*
ns
γ tub+ area (% cell area)
20
15
10
5
0
C
Homeo
Pro-Inf
Anti-Inf
γ tub
GAPDH
48 KDa
37 KDa
Relative o.d. (vs homeo)
1.5
1.0
0.5
0.0
ns
*
Homeo
Pro-inf
Anti-inf
D
Fixation in ice-cold MetOH without extraction
60'
15'
120'
NOC 2 µM 37°C x 1h
Washout and recovery in conditioned medium
Extraction and fixation in ice-cold MetOH
Extraction and fixation in ice-cold MetOH
E
Cx3cr1gfp/+ mice
GFP
IL
OL
F
3D rendering
10 µm
GM130
GFP
G
n° of Golgi outposts
25
20
15
10
5
0
0
10
20
30
40
50
Distance from center (µm)

**Fig. S3**. **Molecular, morphological and functional characterization of microglia non-centrosomal MTs nucleation in primary cultures and in retinal slices**

**(A**) Representative volumetric rendering of g-tubulin (γ tub) signal intensity of homeostatic (Homeo), pro-inflammatory (Pro-inf) and anti-inflammatory (Anti-inf) microglia. **(B)** Scatter dot plot showing analysis of γ tub signal over the cell area in Homeo, Pro-inf and Anti-inf microglia. Values are expressed as mean ± SEM (Homeo n = 17, Pro-inf n = 11 and Anti-inf n = 10 cells from 4 independent experiments). * $p < 0.05$, Kruskal-Wallis test - Dunn's multiple comparison test. **(C)** Representative immunoblot of total γ tub in Homeo, Pro-inf and Anti-inf microglia (*top*); bar chart showing the quantification of γ tub protein levels (*bottom*). Values are expressed as median ± interquartile range from 4 independent experiments. * $p < 0.05$, Mann Whitney test. **(D)** Treatment timeline of Nocodazole wash out assay. **(E)** Representative images of retinal slices (50 μm thickness) from control cx3cr1$^{gfp/+}$ mice stained with Hoechst for nuclei visualization (blue), showing retinal cell layers (IL inner layer, OL outer layer). Scale bar: 20 μm. **(F)** 3D rendering of sample retinal GFP+ microglia (cyan) stained for GM130 (magenta). Scale bar: 10 μm. **(G)** Bar chart reporting the number of Golgi outposts at increasing distance from the center of cell body in retinal microglia.

**Fig.S4**

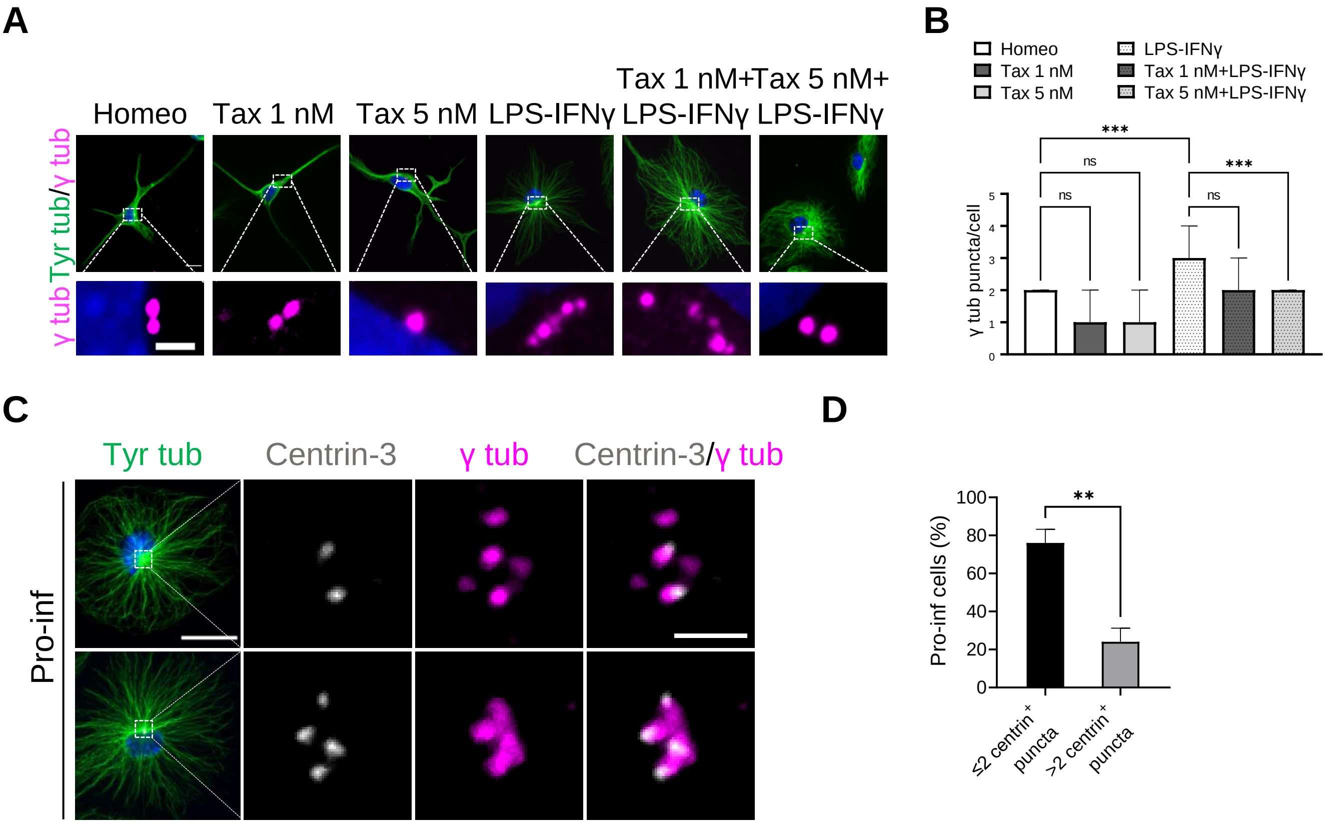


E

| Cell cycle | Homeo | Pro-Inf |
|---|---|---|
| G1 | 83 ± 2 | 75 ± 4 |
| S | 8 ± 1 | 15 ± 2 |
| G2/M | 4 ± 1 | 8 ± 2 |

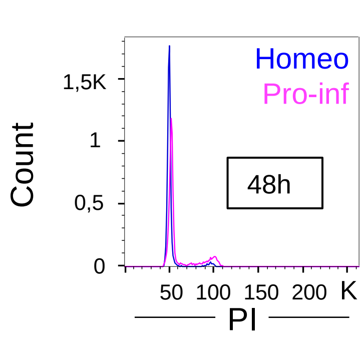


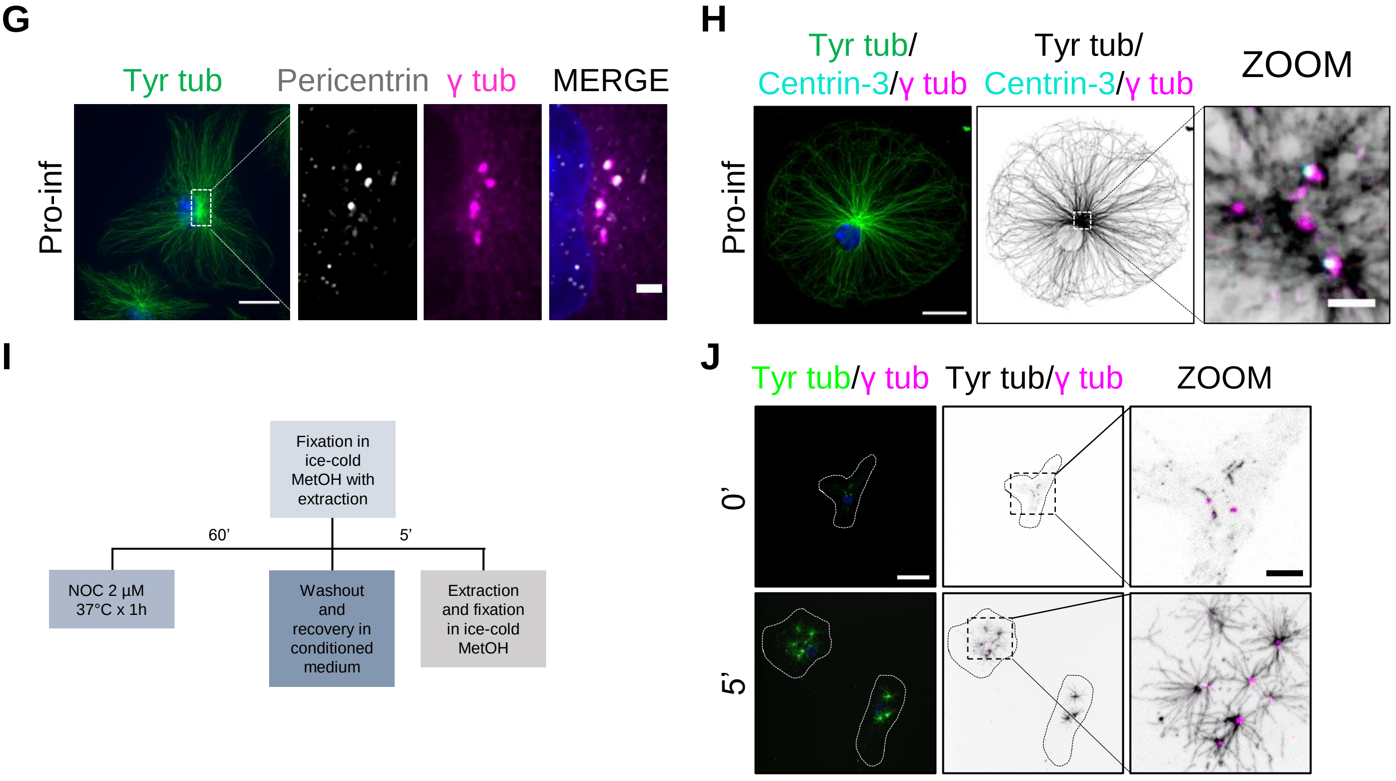

**Fig. S4. Analysis of pericentriolar material maturation during microglia pro-inflammatory activation**

**(A)** Representative images of immunostaining of tyrosinated (Tyr) tub (green) and g-tubulin (γ tub) (magenta) in homeostatic (Homeo), Taxol 1 nM treated (Tax 1 nM), Taxol 5 nM treated (Tax 5 nM), pro-inflammatory (LPS-IFNγ), Taxol 1 nM+LPS-IFNγ treated (Tax 1 nM+LPS-IFNγ) and Taxol 5 nM+LPS-IFNγ treated (Tax 5 nM+LPS-IFNγ) microglia (Top, Scale bar: 10 µm; zoom, 2 µm. Hoechst for nuclei visualization, blue). **(B)** Bar chart reporting the number of γ tub puncta per cell in homeostatic (Homeo), Taxol 1 nM treated (Tax 1 nM), Taxol 5 nM treated (Tax 5nM), pro-inflammatory (LPS-IFNγ), Taxol 1 nM+LPS-IFNγ treated (Tax 1 nM+LPS-IFNγ) and Taxol 5 nM+LPS-IFNγ treated (Tax 5nM+LPS-IFNγ) microglia. Values are expressed as median ± interquartile range from 3 independent experiments. ***p <0.001. Kruskall-Wallis - Dunn's multiple comparisons test. **(C)** Representative images showing Tyr tub (green), centrin-3 (gray) and γ tub (magenta) immunolabeling and co-localization of centrin-3 (gray) and γ tub (magenta) in pro-inflammatory (Pro-inf) microglia. Scale bar: 20 µm; zoom, 2 µm. Hoechst for nuclei visualization, blue. **(D)** Bar graph reporting the percentage of Pro-inf cells displaying ≤2 or >2 centrin$^{+}$ puncta. Values are expressed as mean ± SEM of n = 52 cells from 3 independent experiments. **p <0.01, Student’s t-test. **(E)** Table reporting the percentage of cells in G1, S and G2/M phases from Homeo and Pro-inf microglia cultures, stained with propidium iodide (PI) and analyzed by flow cytometry. Percentages indicate the relative enrichment in cell population, values are expressed as mean ± SEM from three independent experiments. **(F)** Representative histogram of cell cycle overlay of Homeo (blue line) and Pro-inf (magenta line) microglia. **(G)** Representative images showing Tyr tub (green), pericentrin (gray) and γ tub (magenta) immunolabeling in Pro-inf microglia. Scale bar: 20 µm; zoom, 2 µm. Hoechst for nuclei visualization, blue. **(H)** *Left:* representative image of Tyr tub (green), centrin-3 (cyan) and γ tub (magenta) immunolabeling in Pro-inf microglia. *Middle* and *right:* representative image showing Tyr tub (black, inverted LUT), centrin-3 (cyan) and γ tub (magenta) immunolabeling in Pro-inf microglia. Out of focus blur was removed using “remove haze” filter in Metamorph Software to highlight the asters. Scale bar: 20 µm; zoom, 2 µm. Hoechst for nuclei visualization, blue. **(I)** Treatment timeline of Nocodazole wash out assay. **(J)** Representative confocal images of the time course of the MT re-nucleation assay after nocodazole washout in Pro-inf microglia stained for Tyr tub (green) and γ tub (magenta). Scale bar: 20 µm; zoom: 5 µm. Hoechst for nuclei visualization, blue. Time 0’ represents the MT depolymerizing effect of nocodazole in Pro-inf cells with free tubulin extraction.

**Fig.S5**

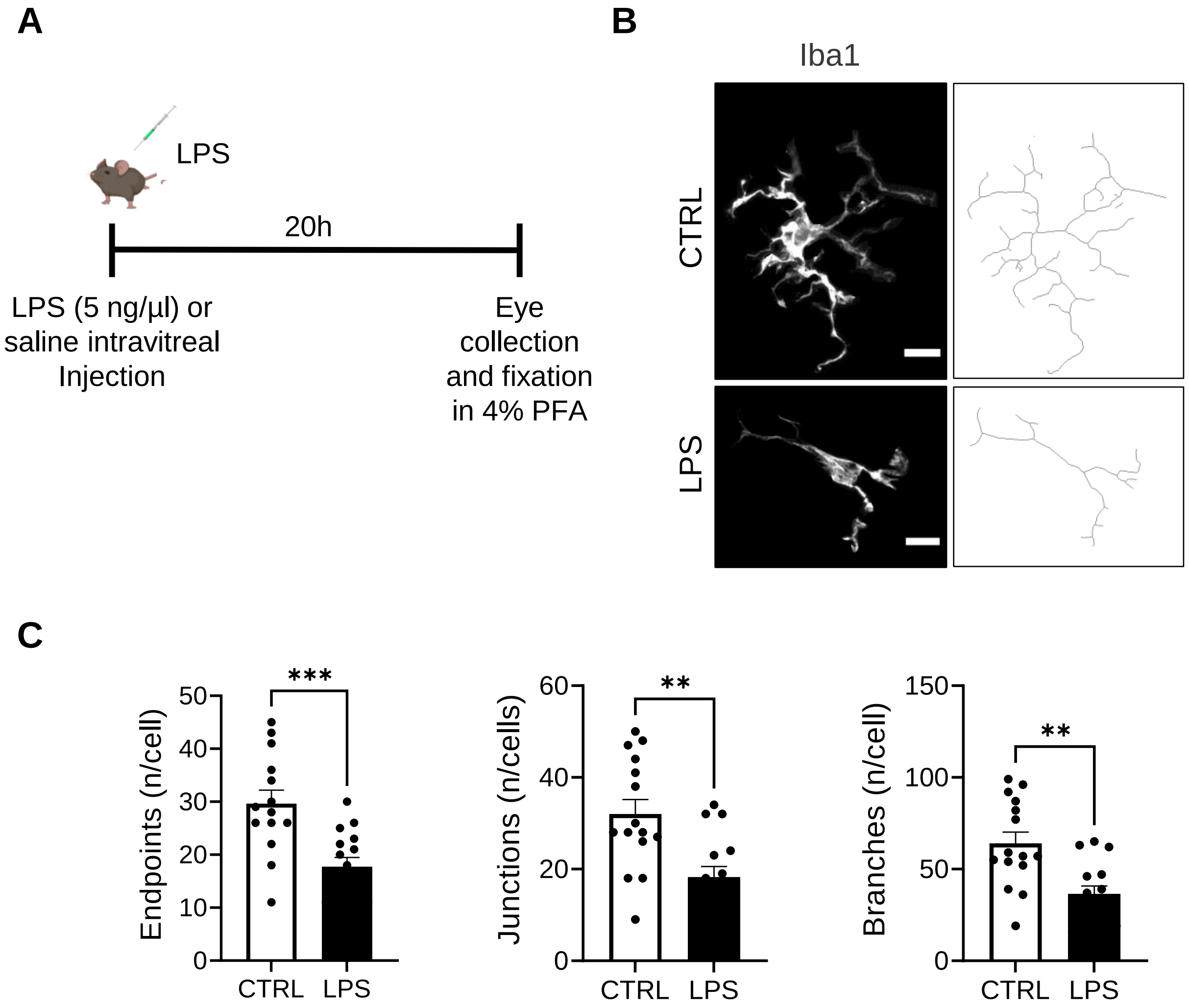

A
LPS
20h
LPS (5 ng/µl) or saline intravitreal Injection
Eye collection and fixation in 4% PFA
B
Iba1
CTRL
LPS
C
Endpoints (n/cell)
Junctions (n/cells)
Branches (n/cell)
***
**
**
CTRL
LPS

**Fig. S5. Characterization of *in vivo* microglia activation in the LPS-induced uveitis model**

**(A)** Schematic illustration of surgical procedure and tissue collection. **(B)** *Left*: representative immunofluorescence images of microglia (Iba1, gray) in retinal slices (50 µm thickness) from CTRL (sham) and LPS treated mice. Scale bar: 10 µm. *Right*: corresponding skeletonized images. **(C)** Scatter dot plots reporting microglia arborization parameters as endpoints (*left*), junctions (*middle*) and branches (*right*) obtained from skeleton analysis of retinal microglia from CTRL (sham) and LPS treated mice. Values are expressed as mean ± SEM (CTRL, n = 14/3 cells/mice; LPS, n = 15/3 cells/mice; *** $p < 0.001$, ** $p < 0.01$; Student's t-test.

**Fig S6**

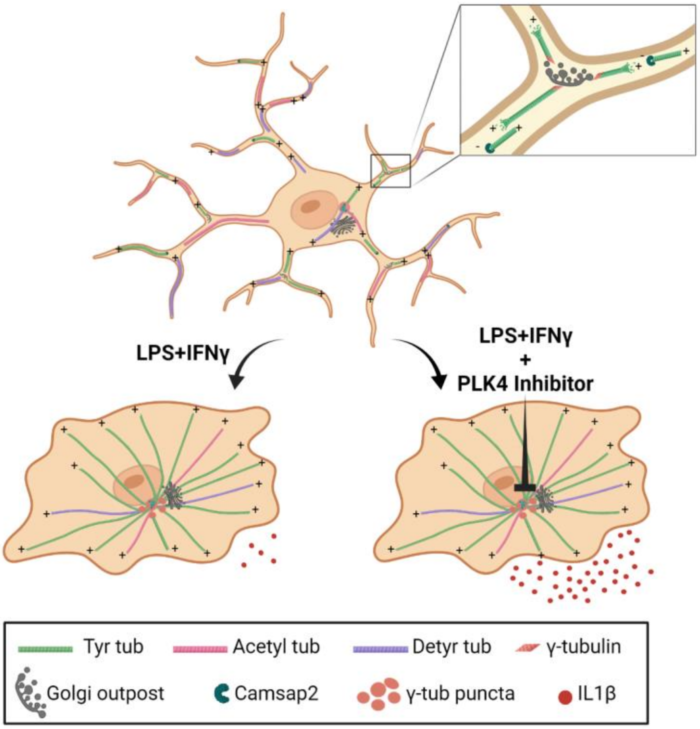
LPS+IFNγ
LPS+IFNγ
+
PLK4 Inhibitor
Tyr tub
Acetyl tub
Detyr tub
γ-tubulin
Golgi outpost
Camsap2
γ-tub puncta
IL1β

**Fig. S6. Graphical abstract to illustrate the main findings of this study.** Homeostatic microglia have stable MT arrays, while microglia reactivity increases MT dynamic behavior. In addition, homeostatic microglia have non-centrosomal MTs with mixed polarity similar to the architecture of highly specialized cells such as neurons and oligodendrocytes. Pro-inflammatory microglia reactivity results in restricted g-tubulin localization to puncta around the centrosome because of *de novo* pericentriolar material (PCM) maturation. Inhibition of PCM maturation by Polo-like Kinase 4 (PLK4) in pro-inflammatory microglia stimulates IL-1b release.

**Fig S7**

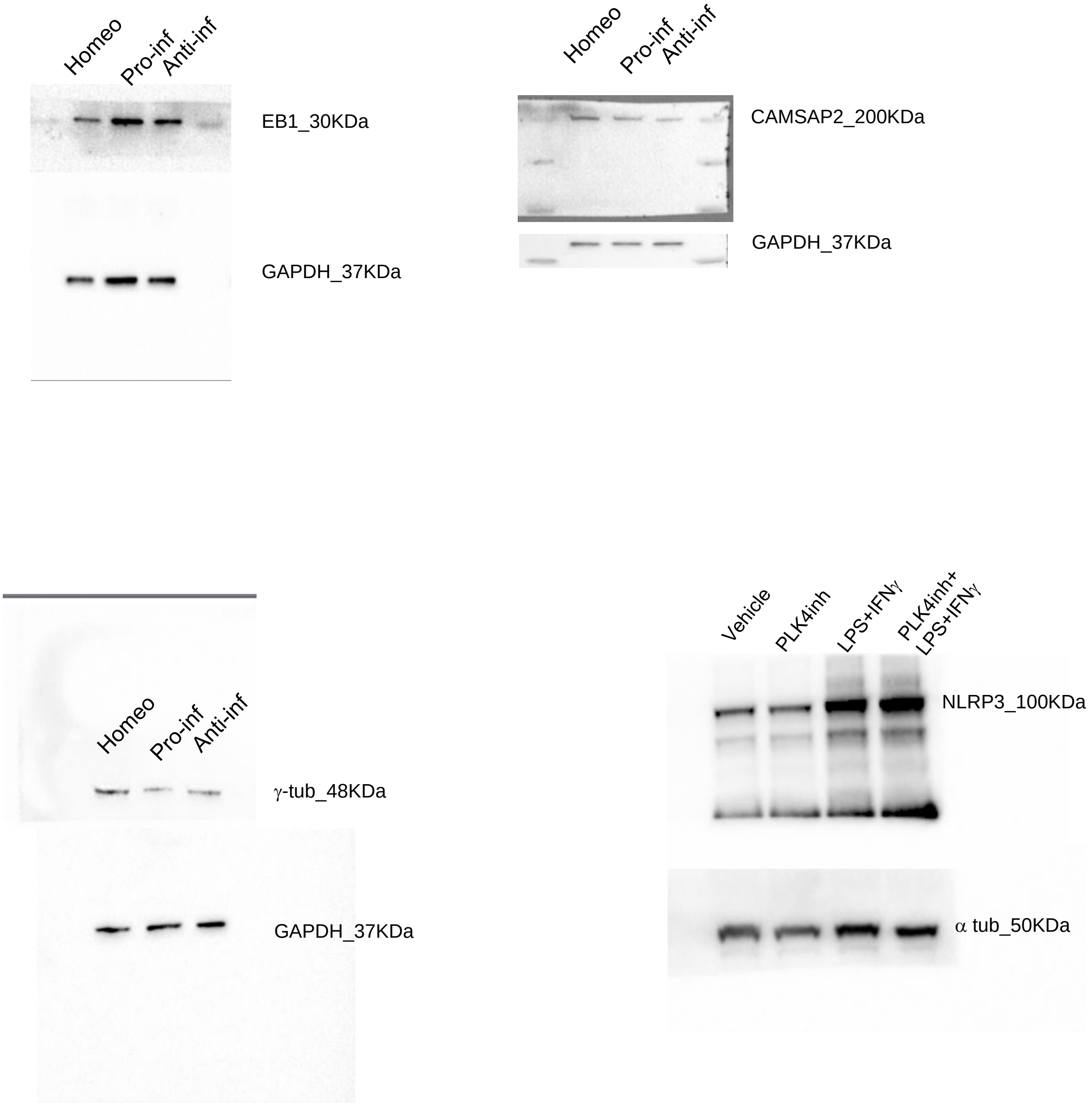


**Fig S7**: Original western blots
